# Expansion of a frontostriatal salience network in individuals with depression

**DOI:** 10.1101/2023.08.09.551651

**Authors:** Charles J. Lynch, Immanuel Elbau, Tommy Ng, Aliza Ayaz, Shasha Zhu, Nicola Manfredi, Megan Johnson, Danielle Wolk, Jonathan D. Power, Evan M. Gordon, Kendrick Kay, Amy Aloysi, Stefano Moia, Cesar Caballero-Gaudes, Lindsay W. Victoria, Nili Solomonov, Eric Goldwaser, Benjamin Zebley, Logan Grosenick, Jonathan Downar, Fidel Vila-Rodriguez, Zafiris J. Daskalakis, Daniel M. Blumberger, Nolan Williams, Faith M. Gunning, Conor Liston

## Abstract

Hundreds of neuroimaging studies spanning two decades have revealed differences in brain structure and functional connectivity in depression, but with modest effect sizes, complicating efforts to derive mechanistic pathophysiologic insights or develop biomarkers.^1^ Furthermore, although depression is a fundamentally episodic condition, few neuroimaging studies have taken a longitudinal approach, which is critical for understanding cause and effect and delineating mechanisms that drive mood state transitions over time. The emerging field of precision functional mapping using densely-sampled longitudinal neuroimaging data has revealed unexpected, functionally meaningful individual differences in brain network topology in healthy individuals,^2–5^ but these approaches have never been applied to individuals with depression. Here, using precision functional mapping techniques and 11 datasets comprising n=187 repeatedly sampled individuals and >21,000 minutes of fMRI data, we show that the frontostriatal salience network is expanded two-fold in most individuals with depression. This effect was replicable in multiple samples, including large-scale, group-average data (N=1,231 subjects), and caused primarily by network border shifts affecting specific functional systems, with three distinct modes of encroachment occurring in different individuals. Salience network expansion was unexpectedly stable over time, unaffected by changes in mood state, and detectable in children before the subsequent onset of depressive symptoms in adolescence. Longitudinal analyses of individuals scanned up to 62 times over 1.5 years identified connectivity changes in specific frontostriatal circuits that tracked fluctuations in specific symptom domains and predicted future anhedonia symptoms before they emerged. Together, these findings identify a stable trait-like brain network topology that may confer risk for depression and mood-state dependent connectivity changes in frontostriatal circuits that predict the emergence and remission of depressive symptoms over time.

---

Depression is a heterogeneous and episodic neuropsychiatric syndrome associated with synapse loss ^6, 7^ and connectivity alterations in frontostriatal networks ^8–10^, and a leading cause of disability worldwide.^11^ The neurobiological mechanisms that give rise to dysfunction in specific depressive behavioral domains or to changes in mood over time are not well understood, especially at the neural systems level. To date, most functional magnetic resonance imaging (fMRI) studies have tested for differences in functional connectivity in cross-sectional comparisons between groups of depressed individuals and healthy, never-depressed controls using group-average (“one-size-fits-all”) parcellations to define functional brain areas and networks. More recently, pioneering work in other areas of systems neuroscience has given rise to the emerging field of precision functional mapping, which refers to a suite of new approaches for delineating functional networks within individuals ^2, 12–19^. Precision mapping studies have shown that the topology (size, shape, spatial location) of functional areas and networks in individuals deviates markedly from group-average descriptions ^2, 4, 20, 21^, and that individual differences in network topology are stable,^2, 20, 22, 23^ heritable,^5, 24^ and associated with cognitive abilities and behavior ^3–5, 13, 25–27^. With the exception of a recent case study involving a single individual who sustained bilateral perinatal strokes^28^, these tools have not yet been widely applied in clinical populations, including depression. Thus, whether functional network topology differs in individuals with depression is unknown.

Importantly, depression is a fundamentally episodic neuropsychiatric condition defined by discrete periods of low mood interposed between periods of euthymia, but our understanding of the mechanisms that mediate mood transitions over time is limited. This is due in part to the fact that most studies to date have been cross-sectional, involving data acquired at a single time point, or in some cases, two or three scans acquired before and after an intervention ^29–32—an^ approach that is not designed for meaningful statistical inferences at the individual level. Understanding the neurobiological mechanisms that mediate transitions in and out of depressive mood states requires dense-sampling of individual patients over many months^33^. Indeed, densely sampled n-of-1 studies involving intracranial EEG recordings and other assessments have begun to reveal mechanisms that regulate mood state transitions in individual patients receiving deep brain stimulation for depression ^34–38^, but these approaches have not yet been deployed at scale in fMRI studies. Absent such datasets, it is unknown whether changes in brain network connectivity predict the emergence of anhedonia, anxiety, and dysfunction in other depressive symptom domains, or the subsequent remission of these symptoms after a recovery from an episode. In the same way, it is unclear whether atypical network topology measures fluctuate with mood state in individuals with depression or remain stable over time—key questions for understanding cause-and-effect relationships in clinical neuroimaging, for defining potential therapeutic targets in neuromodulation interventions, and for identifying biomarkers.

Until recently, technical limitations have posed significant obstacles to performing precision functional mapping and longitudinal neuroimaging in clinical samples, including depression. Conventional fMRI measurements at the single subject level are noisy and have limited reliability, in part because they are sensitive to a variety of imaging artifacts^39^. However, recent studies have developed solutions to these problems, which can be resolved by either acquiring large quantities of data in each subject ^2, 20^ or by using multi-echo fMRI pulse sequences ^23, 40, 41^. Together, these approaches can generate highly reliable functional connectivity measures and network maps at the level of individual subjects, an important first step towards developing and deploying fMRI for clinical translational purposes.

Here, we used state-of-the-art precision functional mapping tools to delineate functional brain network topology in individuals with depression, leveraging multiple datasets comprising >21,000 minutes of resting-state fMRI data. We show that the frontostriatal salience network is expanded by two-fold in most individuals with depression—an effect we replicated in two independent datasets of repeatedly-sampled individuals (n = 106) and in large-scale group-average data (n = 299 depressed patients, N=932 healthy controls), with three distinct types of encroachment on neighboring functional systems occurring across individuals. Salience network expansion was stable over time, and unaffected by changes in mood state. It was also detectable in children scanned before the onset of depression symptoms that emerged later in adolescence. We went on to show that salience network expansion is a highly stable, trait-like feature of individuals at risk for depression, a longitudinal analysis of densely-sampled individuals revealed mood state-dependent changes in striatal connectivity with anterior cingulate and anterior insular nodes of the salience network that tracked fluctuations in anhedonia and anxiety, respectively, and predicted the subsequent emergence of anhedonic symptoms at future study visits.

## Frontostriatal salience network expansion in highly-sampled individuals with depression

Numerous neuroimaging studies involving large cohorts of patients with depression have identified differences in functional connectivity and brain structure ^42–46^, often involving the anterior cingulate cortex, orbitofrontal cortex, insular cortex, and subgenual cingulate cortex—a therapeutic target for deep brain stimulation ^47, 48^—but the effect sizes in large-scale meta-analyses are modest (e.g. Cohen’s *d* = 0.1–0.15 for structural measures ^45^ and *d* = 0.13–0.26 for functional connectivity measures^1^). Whether the topological features of large-scale functional brain networks—their shape, spatial location, and size—are altered in depression is unknown.

We used precision functional mapping to delineate the topology of functional brain networks in six highly sampled individuals with unipolar major depression who underwent on average 621.5 minutes of multi-echo fMRI scanning (range: 58–1,792 minutes) across 22 sessions (range: 2– 62 sessions). We refer to this dataset as the Serial Imaging of Major Depression (SIMD) dataset (see study design and aims in **Extended Data** Fig. 1a). To contextualize the severity of depressive symptoms in these individuals, the mean 17-item ^49^ Hamilton Depression Rating Scale (HDRS17) score (averaged across study visits, excluding those when these individuals were in remission) was 15.7 ± 3.7 (range: 10.5–22.2), indicating a range of severity levels from mild to severe. The same precision mapping procedures were applied to 36 highly-sampled healthy controls with an average of 317.3 minutes of fMRI data per subject (range: 43.0–841.2 minutes) across 12 sessions (range: 2–84 sessions). See the Methods section for additional details. The size of each functional network was quantified in each individual as the proportion of the total cortical surface area it occupied (see **Extended Data** Fig. 1b and Methods section for additional details).

**Fig. 1.**
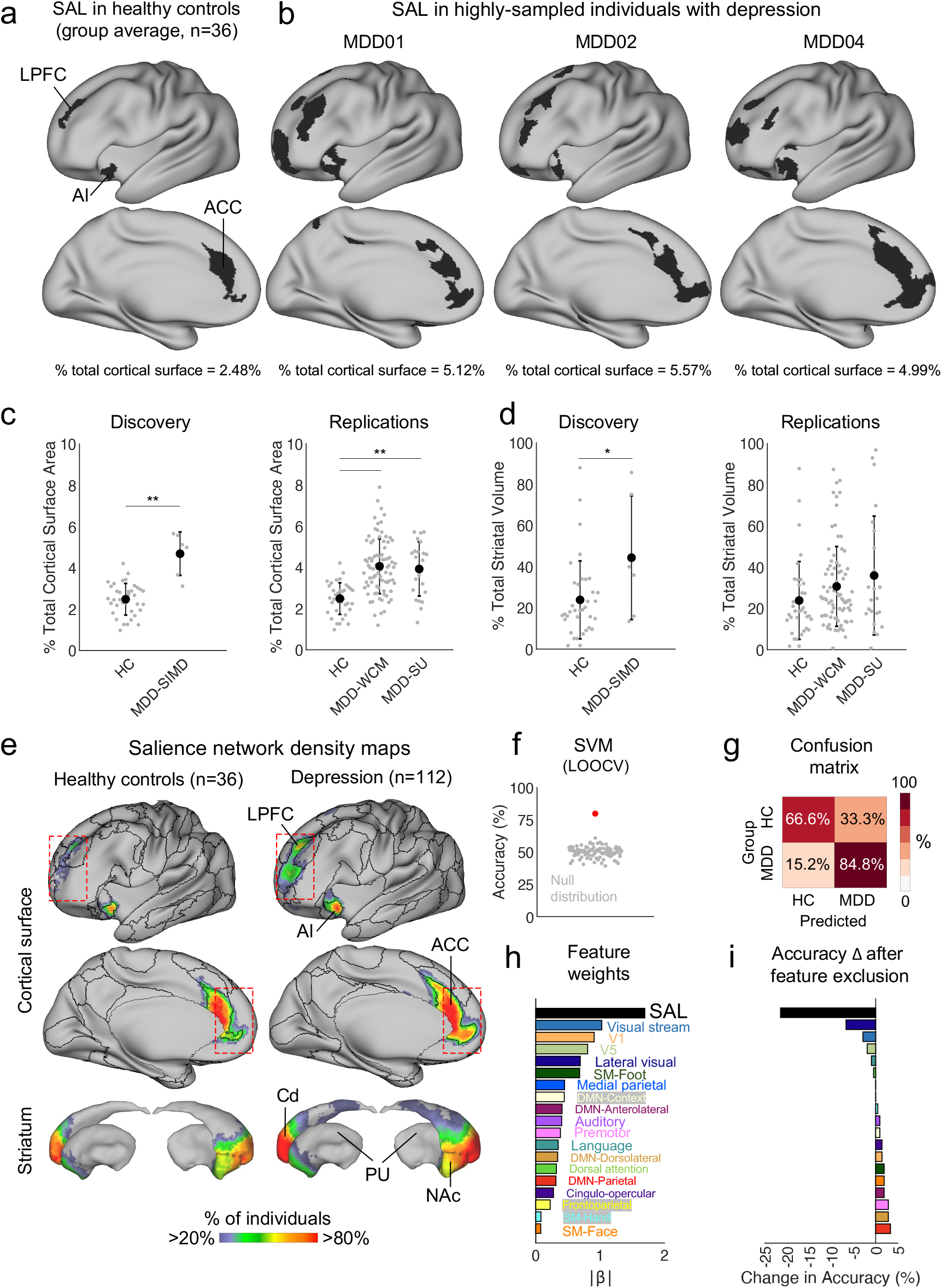
Frontostriatal salience network is expanded 2-fold in highly-sampled individuals with depression. **a**, The salience network (SAL, black) has cortical representation in lateral prefrontal cortex (LPFC), anterior cingulate cortex (ACC), and anterior insular cortex (AI), and occupied 2.48% of the cortical surface on average in n = 36 highly-sampled healthy controls (HC). **b-c,** The salience network was approximately 2x larger in n = 6 highly-sampled individuals with depression (a dataset referred to here as Serial Imaging of Major Depression, MDD-SIMD), occupying 88.9% more of the cortical surface on average (permutation test, \*\**P* = 0.001, Bonferroni correction, *Z*-score = 6.20). This effect was replicated twice using data from separate imaging sites (two-tailed independent sample *t*-tests, n = 83 from Weill Cornell Medicine, MDD-WCM: *t* = 8.09, \*\**P* < 0.001, Bonferroni correction, Cohen’s *d* = 1.12; n = 23 from Stanford University, MDD-SU: *t* = 4.75, \*\**P* < 0.001, Bonferroni correction, Cohen’s *d* = 1.16). **d,** The total striatal representation of the salience network was also increased relative to healthy controls (permutation test, \**P* = 0.02, uncorrected, *Z*-score = 2.34), although this effect was not statistically significant in either replication dataset (two-tailed independent sample *t*-tests, MDD-WCM: *t* = 1.78, *P* = 0.07, uncorrected; MDD-SU: *t* = 1.79, *P* = 0.08, uncorrected). **e,** Density maps indicated that the spatial locations of salience network nodes were similar in healthy controls and individuals with depression, but that network borders tended to extend further outwards from their centroids in each cortical zone in depression (see red boxes). **f-i**, A support vector machine (SVM) classifier trained using leave-one-out cross-validation (LOOCV) distinguished depressed individuals from healthy controls (accuracy = 80%, significance assessed using a permutation test, *P* = 0.001) using only the size of each functional network as features. Linear predictor coefficients (β) associated with the trained model and the change in accuracy after feature exclusion both indicated that salience network size was the most important feature. Error bars represent standard deviation.

It was immediately apparent upon visual inspection that the salience network, which is involved in reward processing and conscious integration of autonomic feedback and responses with internal goals and environmental demands ^50, 51^, was markedly expanded in these individuals with depression (**Fig. 1a-b**). In four of the six individuals, the salience network was expanded more than two-fold, outside the range observed in all 36 healthy controls (**Fig. 1c**, left). On average, the salience network occupied 88.7% more of the cortical surface relative to the mean in healthy controls (4.68% ± 1.05% of cortex in SIMD versus 2.48% ± 0.74% of cortex in healthy controls), giving rise to a large group-level effect (Cohen’s *d* = 1.97). To further validate this observation, we repeated this procedure in two additional datasets (n = 83 from Weill Cornell Medicine and n = 23 from Stanford University) of individuals with depression repeatedly sampled using multi-echo fMRI (58 to 86 minutes of data per subject). The effect was replicated in both samples (**Fig. 1c**, right), again with large effect sizes (Cohen’s *d* = 1.12 – 1.16), and remained statistically significant when controlling for the sex ratio imbalance in our samples (58% of individuals with depression were female, versus 31% of healthy controls, see **Extended Data** Fig. 2). We also found greater subcortical representation of the salience network in the striatum (**Fig. 1d**, left), which is thought to relate to anatomically well defined, interconnected loops in which the cortex projects to the striatum, and the striatum projects back to cortex indirectly via the thalamus ^52, 53^.

**Fig. 2.**
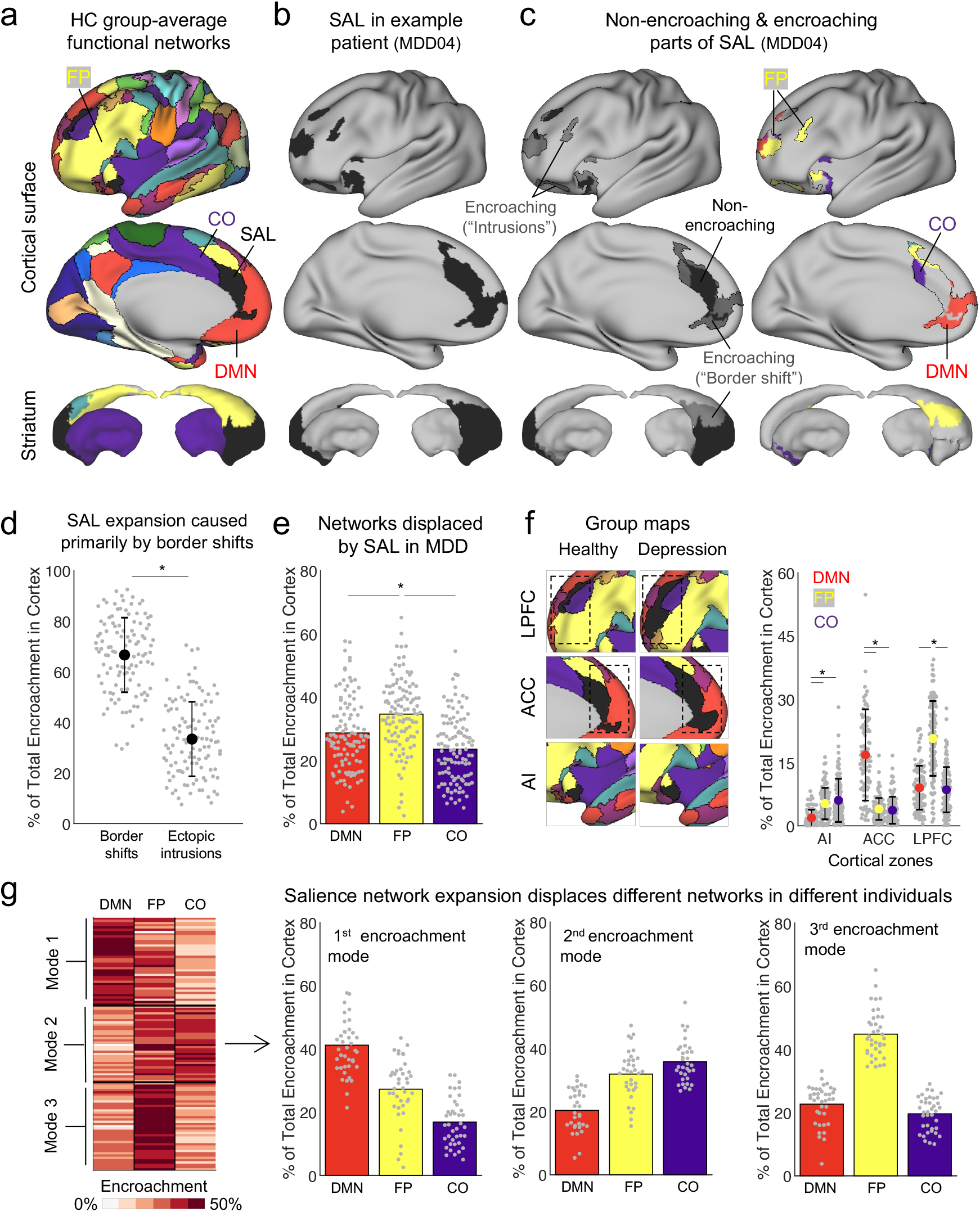
Three modes of network encroachment underlying salience network expansion in depression. **a**, Functional brain networks, including the salience network (SAL, black), mapped using group-average healthy control (HC) data. **b-c**, The parts of each depressed individual’s salience network that did and did not overlap with the healthy controls are referred to as non-encroaching and encroaching, respectively. **d**, The relative contribution of each functional network to the total surface area of the encroaching portion of the salience network was calculated to quantify which functional networks were encroached upon most, referred to as their encroachment profile. Salience network expansion was primarily caused by network borders shifts, and to a lesser extent ectopic intrusions – isolated patches of salience network in atypical locations (two-tailed paired sample *t*-test, *t* = 11.94, \**P* < 0.001). **e**, Three functional systems, the default mode (DMN, red), frontoparietal (FP, yellow) and cingulo-opercular (CO, purple) networks were most affected by salience network expansion. When considering the entire cortical surface, the frontoparietal network was more affected compared to the default mode (two-tailed paired sample *t*-test, *t* = 2.89, \**P* = 0.004) and cingulo-opercular (two-tailed paired sample *t*-test, *t* = 6.92, \**P* < 0.001) networks. **f**, However, the functional network was displaced most by salience network expansion differed by cortical zone. In the anterior insular cortex (AI), the frontoparietal network (two-tailed paired sample *t*-test, *t* = 7.39, \**P* < 0.001, Bonferroni correction) and cingulo-opercular (two-tailed paired sample *t*-test, *t* = 6.92, \**P* < 0.001, Bonferroni correction) networks were more affected than the default mode network. In the anterior cingulate cortex (ACC), the default mode network was more affected than either the frontoparietal (two-tailed paired sample *t*-test, *t* = 12.01, \**P* < 0.001, Bonferroni correction) or cingulo-opercular (two-tailed paired sample *t*-test, *t* = 11.71, \**P* < 0.001, Bonferroni correction) networks. Finally, in lateral prefrontal cortex (LPFC), the frontoparietal network was more affected than either the default mode (two-tailed paired sample *t*-test, *t* = 11.83, \**P* < 0.001, Bonferroni correction) or cingulo-opercular (two-tailed paired sample *t*-test, *t* = 12.21, \**P* < 0.001, Bonferroni correction). **g**, Clustering individuals (using the Louvain method of community detection) using their encroachment profiles revealed three distinct modes of encroachment across individuals. Error bars indicate SD.

Salience network expansion in depression was also evident in density maps (**Fig. 1e**), which convey the percentage of individuals with salience network representation at each cortical vertex or striatal voxel. These maps confirmed a similar overall pattern of cortical and subcortical representation in both groups, consistent with descriptions in previous reports ^2, 51, 54^, but also revealed that the borders of the salience network tended to extend further outwards from their centroids in each cortical zone in depressed individuals. For example, in the anterior cingulate cortex, network borders shifted more anteriorly into the pregenual cortex, and in lateral prefrontal cortex, network borders shifted more anteriorly towards the frontal pole (see red boxes in **Fig. 1e**). Accordingly, expansion of the salience network in cortex and striatum was accompanied by contraction of three neighboring functional systems: the default mode, frontoparietal, and cingulo-opercular systems (**Extended Data** Fig. 3). However, the specific patterns of contraction did not replicate in all datasets—a finding we return to in the following section. Otherwise, consistent and reproducible group differences in network size were specific to the salience network: while the frontotemporal language network was also larger than normal in the discovery sample (MDD-SIMD: permutation test, *P* = 0.011, Bonferroni correction, *Z*-score = 4.61), this effect was reproducible in only one of the replication samples (two-tailed independent sample *t*-tests, MDD-WCM: *t* =0.50, *P* = 0.62, uncorrected, Cohen’s *d* = 0.10; MDD-SU: *t* = 4.25, *P* = 0.004, Bonferroni correction, Cohen’s *d* = 0.99), and there were no significant differences in the size of any other network after correcting for multiple comparisons.

**Fig. 3.**
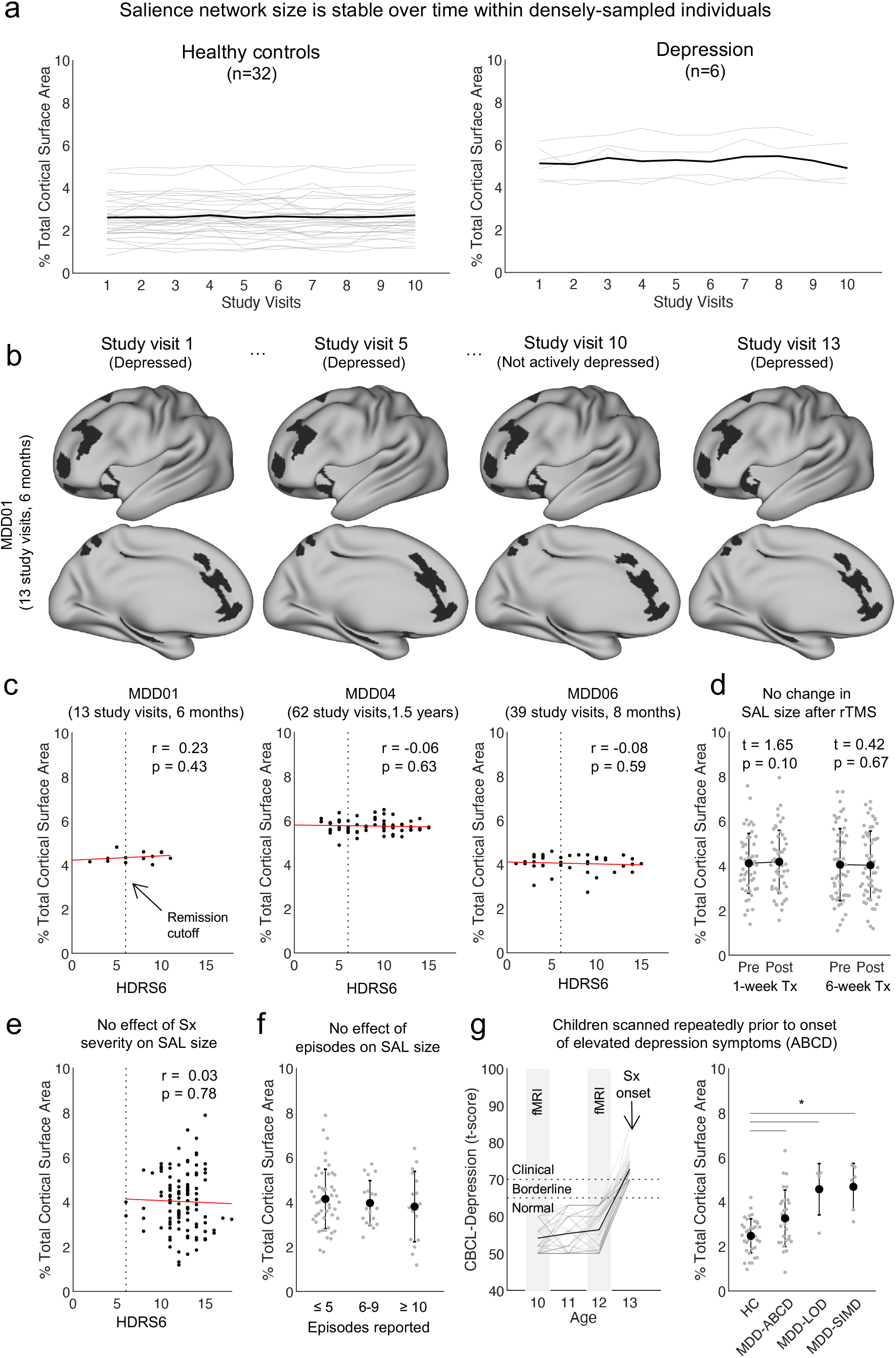
Salience network expansion is stable over time and emerges early in life. **a**, The overall cortical representation of the salience network is highly stable in repeatedly scanned healthy controls (left) and individuals with depression (right). The first ten study visits for each subject are shown for visualization purposes. **b**, Salience network topology across study visits and mood states in a representative individual with depression scanned longitudinally. **c-d,** No significant correlation between the overall severity of depressive symptoms (Hamilton Depression Rating Scale, HDRS6) and salience network size in any repeatedly-sampled individual with depression (Pearson correlation, all p’s > 0.4), or in patients before and after a course of either an accelerated 1-week (two-tailed paired sample *t*-test, *t* = 1.65, *P* = 0.10, uncorrected, n = 50) or traditional 6-week (two-tailed paired sample *t*-test, *t* = 0.42, *P* = 0.67, uncorrected, n = 53) repetitive transcranial magnetic stimulation therapy (rTMS). **e-f**, Cross-sectional analysis using all depression subjects (n = 112) revealed that individual differences in salience network size are not related to the overall severity of symptoms (HDRS6, Pearson correlation, *r* = 0.03, *P* = 0.78, uncorrected), nor with the number of depressive episodes individuals reported having during their lifetime (inferred from the Mini-International Neuropsychiatric Interview). **g**, A subset (n=54) of children from the Adolescent Brain Cognitive Development (ABCD) study scanned prior to the onset of elevated depression symptoms were identified. Depression symptoms were operationalized using the DSM-oriented scale for depression from the Child Behavior Checklist (CBCL, *T*-scores ≥ 70 are in the clinical range). The salience network was significantly larger in children who later developed depression (MDD-ABCD, permutation test, \**P* = 0.002, uncorrected, *Z*-score = 3.52) and in older adults meeting criteria for late-onset depression (MDD-LOD, permutation test, \**P* = 0.001, uncorrected, Z-score = 5.65) relative to healthy controls (HC, same data also shown in **Fig. 1c**). Sx = symptom, Tx = treatment. Error bars indicate standard deviation.

To better understand whether this effect was also detectable in large, previously published samples involving conventional single-echo fMRI data, we identified the salience network in group-average functional connectivity data from two large datasets involving n = 812 ^55^ and n = 120 ^54^ healthy control subjects, respectively, and in a third dataset involving n = 299 individuals with treatment-resistant depression scanned in association with a neuromodulation intervention study ^56^. The cortical representation of the salience network was >70% larger in the 299-subject depression sample compared to two healthy control samples (**Extended Data** Fig. 4). Furthermore, highly similar patterns of salience network topology and functional connectivity were produced in split-half analyses of each SIMD dataset (**Extended Data** Fig. 5), indicating that salience network expansion was a robust and reproducible feature of these highly-sampled individuals’ brains.

**Fig. 4.**
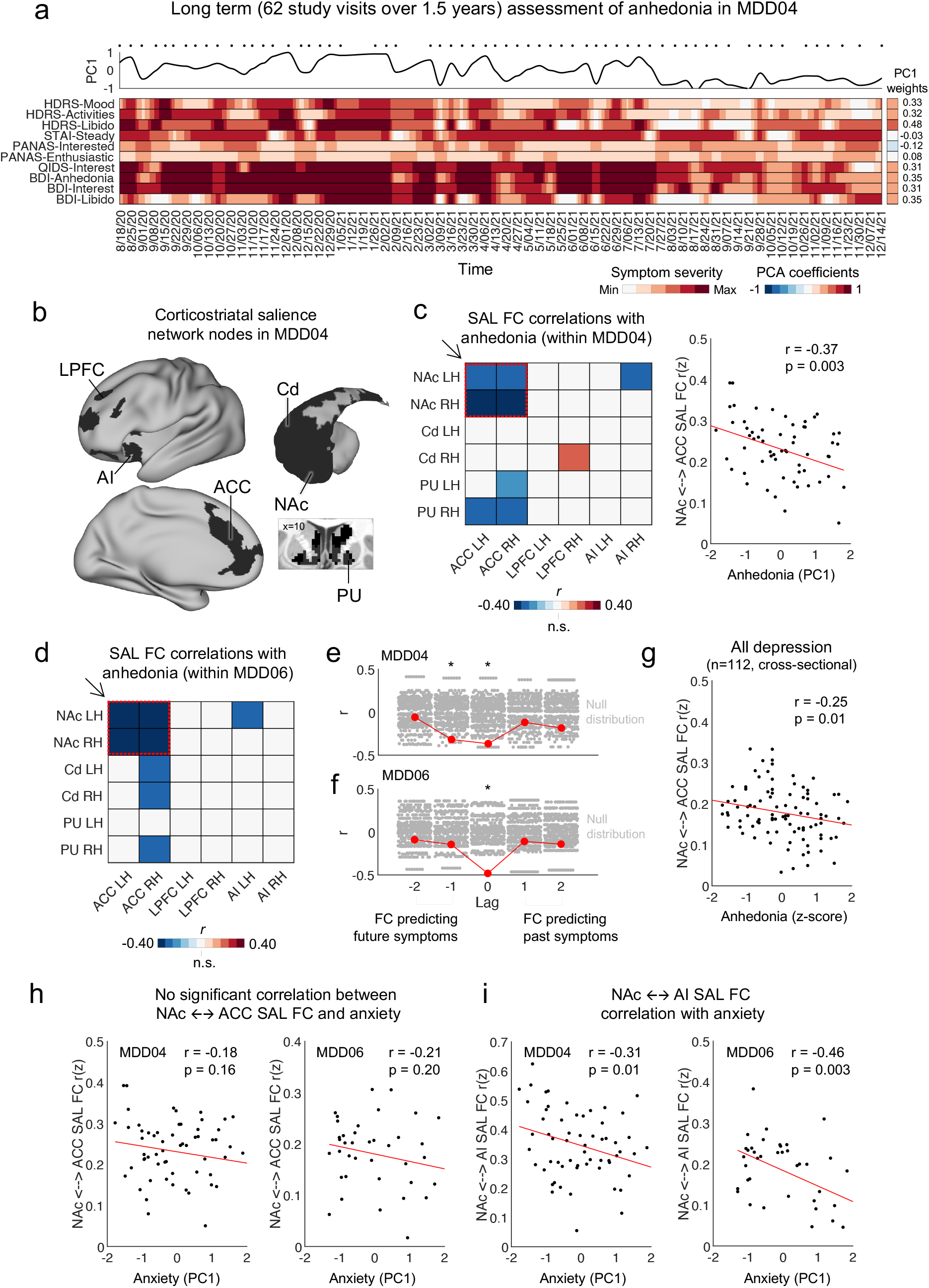
Frontostriatal salience network connectivity predicts fluctuations in anhedonia over time. **a,** A heat map summarizes fluctuations in individual items selected from a variety of clinical interviews and self-report scales related to anhedonia in an example individual (MDD04). Clinical data was resampled (using shape-preserving piecewise cubic interpolation) to days for visualization purposes (black dots above heat map mark the study visits). **b,** The frontostriatal nodes of the salience network in MDD04. **c-d,** Salience network functional connectivity (FC), and notably between nodes in the nucleus accumbens (NAc) and anterior cingulate cortex (ACC), tracked fluctuations in the severity of anhedonia related symptoms within both MDD04 (Pearson correlation, *r* = -0.37, *P* = 0.003) and MDD06 (Pearson correlation, *r* = -0.48, *P* = 0.001) across study visits. FC was calculated as the correlation between the average time courses of all salience network vertices or voxels in a given cortical zone and striatal structure. Statistical significance was assessed using permutation tests with circular rotation to preserve temporal autocorrelation. **e-f**, Cross-correlation analyses indicated NAc ←→ ACC FC also predicted the severity of anhedonia related symptoms at the following study visit in MDD04 (Pearson correlation, significance tested via permutation test, *r* = -0.32, \**P* = 0.004), but not in MDD06. **g**, There was also a significant negative correlation between individual differences in salience network NAc ←→ ACC FC strength and the severity of anhedonia related symptoms (at baseline) when using the entire study sample cross-sectionally (Pearson correlation, *r* = 0.25, *P* = 0.01). **h-i**, Within MDD04 and MDD06, salience network NAc ←→ ACC FC was not significantly related to fluctuations in the severity of other depressive symptoms, such as anxiety, whereas FC between other pairs of nodes (nucleus accumbens and anterior insula) was, indicating specificity with respect to which patterns of functional connectivity relate to different core depressive symptoms (Pearson correlations, MDD04: *r* = -0.31, *P* = 0.01; MDD06: *r* = -0.46, *P* = 0.003).

Given the magnitude of the effect reported in **Fig. 1c** (Cohen’s *d* = 1.12–1.98), we went on to test whether individuals with depression could be distinguished algorithmically from healthy controls using only the size of each functional network (a low-dimensional description of functional brain organization) as predictive features. Although fMRI-based classifiers typically correct for scanner-related artifacts by regressing or co-varying for scanner effects that may influence individual functional connectivity measures, we hypothesized that these corrections may not be necessary here because group differences in salience network size were visually apparent across datasets acquired on different scanners. Thus, we trained a linear support vector machine classifier to differentiate individuals with depression from healthy control individuals based on the size of all 19 functional networks, pooling data from n = 36 healthy controls acquired from 5 different scanners and n = 112 individuals with depression acquired from two different scanners / manufacturers (i.e. all the data in **Fig. 1c**). Classifier performance was evaluated through leave-one-out cross validation, with statistical significance assessed through permutation testing with shuffled diagnostic labels. To prevent classification bias attributable to differences in the size of each group, we oversampled healthy controls using synthetic data derived exclusively from the training sample to prevent data leakage as in ^57^. Overall, support vector machine classifiers correctly differentiated depression cases from healthy controls with 80.4% accuracy (permutation test, *P* = 0.001; **Fig. 1f**), correctly diagnosing depression in 84.8% of cases (**Fig. 1g**), for a positive predictive value of 88.8%. Feature importance was evaluated by examining the linear predictor coefficients and calculating the change in accuracy after exclusion (**Fig. 1h-i**). As expected, salience network size was the most distinguishing feature. Together, these analyses indicate that the salience network is markedly expanded in individuals with depression, with large effect sizes that are reproducible in multiple samples involving different data acquisition and analysis procedures and sufficient in magnitude to support individual classifications with high accuracy rates.

## Three types of network encroachment underlying salience network expansion

Individual differences in functional brain organization are known to occur in two forms: ectopic intrusions, in which isolated pieces of a functional network are observed in an atypical location, and border shifts, in which the boundary of a network expands (or contracts) and encroaches on its neighbors^4^. Border shifts are heritable^5^ and more easily explained by known mechanisms of cortical expansion that are controlled by genetic programs that refine boundaries between functional areas during development and with experience or in response to environmental influences^58^. Furthermore, macroscale networks in both humans and non-human primates are organized in a hierarchy associated with cortical gradients in gene expression and functional properties ^59^, with unimodal sensorimotor areas at the base and heteromodal association areas such as the default mode network at the apex ^60^. Thus, as a first step toward understanding the mechanisms that give rise to cortical expansion of the salience network in depression, we tested whether it was driven primarily by border shifts or ectopic intrusions, and whether it tended to affect lower-level, unimodal sensorimotor networks or heteromodal association areas positioned higher in this hierarchy. To this end, we first generated a central tendency functional network map for the 36 healthy controls (**Fig. 2a**). Second, we identified parts of the salience network in each of the 112 individuals with depression that did not overlap with the salience network in the group-average map for healthy controls, and classified them as either ectopic intrusions or border shifts (**Fig. 2b-c**). Next, we calculated an encroachment profile for each subject, by quantifying the degree of encroachment on every other functional network, defined as the relative contribution of each functional network to the total surface area of the encroaching portion of the salience network (**Fig. 2c**).

This analysis confirmed that salience network expansion was not randomly distributed—instead, it was due primarily to border shifts affecting three neighboring higher-order functional systems, with three distinct encroachment profiles occurring in different individuals. Although salience network expansion involved both ectopic intrusions and border shifts, the latter were more common (**Fig. 2d**), and both tended to result in encroachment on the default, frontoparietal, or cingulo-opercular networks (**Fig. 2e**), not unimodal sensorimotor networks (**Extended Data** Figure 6). It was also evident that the salience network tended to encroach upon specific functional networks in different cortical zones (**Fig. 2f**). For example, in the lateral prefrontal cortex, the salience network expanded rostrally and tended to displace the frontoparietal network. In contrast, in the anterior cingulate and anterior insular cortex, the default mode and cingulo-opercular networks were disproportionately affected, respectively. Clustering individuals by their encroachment profiles revealed three distinct modes (**Fig. 2g**), involving predominantly the default mode network, the frontoparietal network, or a combination of the frontoparietal and cingulo-opercular networks. This heterogeneity may partly explain our observation that the salience network was consistently expanded in all three datasets, but corresponding contractions of other functional networks were more variable.

The results above indicate that salience network expansion is driven primarily by encroachment upon the frontoparietal, cingulo-opercular, and default mode networks, and suggest that cortical space at the boundary between networks may be allocated to different functional systems in individuals with depression. To test this, and to further validate our findings, we compared the strength of functional connectivity between encroaching nodes of the salience network (dark gray vertices in the left part of **Fig. 2c**), and the functional networks that typically occupy that space in healthy controls. As expected, the functional connectivity of encroaching salience network nodes with the rest of the salience network (mean r_z_ = 0.29) was significantly stronger than with displaced networks (all mean r_z_ < 0.03), consistent with a nearly complete disconnection of most encroaching nodes from the functional networks that typically occupy that space in healthy controls (**Extended Data** Fig. 7). Together, these results show that frontostriatal salience network expansion is driven primarily by network border shifts and encroachments that affect three specific higher-order functional systems and spare others, with distinct modes of encroachment occurring in three subgroups of patients.

## Salience network expansion is stable over time and emerges early in life

Major depressive disorder is a fundamentally episodic condition defined by discrete periods of low mood interposed between periods of euthymia ^61, 62^. The salience network has been implicated in reward processing and in integrating autonomic signals with internal goals and changing environmental contexts ^50, 51^. Thus, we hypothesized that changes in salience network topology may precede and predict changes in hedonic function and other symptom domains that occur during mood state transitions—a hypothesis that our longitudinal SIMD dataset was especially well-suited to test. Unexpectedly, we found that salience network topology was highly stable over time, in both healthy controls and individuals with depression (**Fig. 3a-b**). Within-subject analyses revealed no significant correlation between fluctuations in depression symptoms (HDRS6, a more sensitive measure of changes on shorter timescales) and changes in salience network size over time in any of the densely-sampled individuals in our SIMD dataset (**Fig. 3c**). To address the same question, we asked whether salience network size changed after antidepressant treatment, leveraging two cohorts of patients scanned before and after an accelerated, one-week intensive course of repetitive transcranial magnetic stimulation (rTMS; n = 50) or a conventional six-week course of rTMS treatment (n = 56). There were no significant changes in salience network size in either sample (**Fig. 3d**). In addition, neither the severity of symptoms during a given episode (**Fig. 3e**), nor the total number of depressive episodes individuals reported experiencing during their lifetime (**Fig. 3f**) explained individual differences in salience network size. Collectively, these findings indicate that contrary to our initial hypothesis, salience network topology is a stable feature of individuals with major depressive disorder, and unrelated to an individual’s current mood state, to the severity of their symptom severity, or to the chronicity of their illness.

These observations led us to hypothesize that instead of driving mood state-dependent changes in depressive symptoms over time, salience network expansion may be a stable marker of risk for developing depression. To test this hypothesis, we asked whether increased cortical representation of the salience network was detectable early in life, before the onset of depressive symptoms in individuals who would go on to develop depression subsequently. Using data from the Adolescent Brain Cognitive Development (ABCD) study ^63, 64^, we identified n = 54 children who were scanned repeatedly at ages 10 and 12 who did not have significant depressive symptoms in their initial study visits but went on to develop clinically significant depressive symptoms at age 13 (**Fig. 3g**). Precision functional mapping in these individuals revealed that on average, the salience network occupied 38.8% more of the cortical surface in children with no current or prior symptoms of depression at the time of their fMRI scans, but who subsequently developed clinically significant symptoms of depression (**Fig. 3g**: 3.27% ± 1.27% of cortex in MDD-ABCD versus 2.42% ± 0.74% of cortex in healthy control adults, also shown in **Fig. 1c**). A similar effect was also observed in adults with late-onset depression (**Fig. 3g**: MDD-LOD, operationalized as the first episode occurring after age 60). Together, these results show that cortical expansion of the salience network is a trait-like feature of brain network organization that is stable over time, unaffected by mood state, and detectable in children prior to the onset of depression symptoms in adolescence.

## Frontostriatal salience network connectivity predicts fluctuations in anhedonia over time

The results above indicate that topological features of the salience network such as its size, shape, and spatial location are stable over time and do not fluctuate with mood state. However, this observation does not rule out the possibility that functional connectivity between specific salience network nodes may fluctuate in strength, and that such fluctuations could contribute to the emergence of depressive episodes and their subsequent remission. To test this, we first asked whether changes in functional connectivity strength between nodes of the salience network either co-occur with or predict fluctuations in symptom severity over time within individuals, focusing initially on hedonic function, a core feature of depression that has been associated with frontostriatal circuits ^65–70^. Our analyses focused on two of the patients from the SIMD dataset (MDD04 and MDD06) who were repeatedly scanned and assessed by clinicians longitudinally over 8–18 months, providing sufficient data for this analysis. This afforded an opportunity to ask for the first time at the level of single, densely sampled individuals—whose data effectively served as independent, well-powered n-of-1 experiments^18^—how variability in brain network functional connectivity relates to fluctuations in specific symptom domains.

We began with MDD04 because this individual was studied over the longest period of time (62 study visits over 1.5 years) and had the most fMRI data (29.96 hours in total), and reserved MDD06 for replication purposes (39 study visits over 8 months, 18.85 hours in total). During a period spanning over 1.5 years, we observed significant fluctuations in ten anhedonia-related measures (**Fig. 4a**), which were derived from five standardized depressive symptom scales and identified by a consensus clinical decision by three study co-authors (see Methods), ranging from mild / negligible to severe. We tested whether changes in functional connectivity between nodes of the salience network were correlated with changes in anhedonia in this individual over time, as measured by a principal component analysis of the ten anhedonia-related measures in **Fig. 4a** and summarized by the first component score. To reduce the likelihood of false positives, this analysis focused on connectivity between the three major cortical (anterior cingulate, lateral prefrontal, anterior insula cortex) and striatal (nucleus accumbens, caudate, putamen) nodes of the salience network. We found that functional connectivity between multiple cortical and striatal salience network nodes was correlated with changes in anhedonia over time (**Fig. 4b-c**), with the strongest effects observed for connectivity between the nucleus accumbens and anterior cingulate cortex. An identical analysis in MDD06, involving 39 study visits over 8 months, replicated this effect (**Fig. 4d**, **Extended Data** Fig. 8a-b). Bootstrap resampling, a measure of reliability ^71^, indicated good stability for the relationship between anhedonia and nucleus accumbens–anterior cingulate functional connectivity within both individuals (**Extended Data** Fig. 8c). Finally, to determine whether changes in nucleus accumbens–anterior cingulate functional connectivity not only predicted changes in anhedonia within individual subjects but also explained individual differences in anhedonia at a given point in time, we repeated this analysis cross-sectionally in the n = 112 subject replication cohort used in our work above (**Fig. 1c-e**). This analysis confirmed that salience network connectivity between the anterior cingulate and nucleus accumbens was also correlated with anhedonia across individuals (**Fig. 4g**), albeit with a somewhat smaller effect (Pearson’s *r* = –0.25 in the cross-sectional analysis versus Pearson’s *r* = –0.37 and –0.48 for the within-subject analyses), underscoring the value of within-subject analyses.

Finally, we asked whether salience network functional connectivity was predictive of symptom severity at *future* study visits and whether the effect was specific to anhedonia or extended to other symptom domains. Remarkably, a cross-correlation analysis examining correlations with symptoms in past, present, and future study visits showed that functional connectivity between the salience network nodes in the nucleus accumbens and anterior cingulate was not only correlated with current anhedonia symptoms but also predicted the future emergence or remission of anhedonia symptoms in the next study visit in MDD04 (**Fig. 4e**), typically with a lag of approximately one week. The significance of this effect was confirmed using permutation tests with circular rotation to preserve temporal autocorrelation, indicating that accumbens– anterior cingulate connectivity at a given visit predicted future anhedonia approximately one week later, even after controlling for correlations in anhedonia measures over time. Of note, salience network connectivity correlations were replicated in MDD06 for current symptoms, but not for future symptoms (**Fig. 4f**), which may relate to differences in their antidepressant treatments in this observational setting (MDD06 was undergoing maintenance electroconvulsive treatment unrelated to this study).

To evaluate the specificity of this effect, we asked whether nucleus accumbens–anterior cingulate connectivity was also correlated with fluctuations in anxiety symptoms over time and observed no significant correlation in either individual (**Fig. 4h**), indicating a specific role for accumbens–anterior cingulate connectivity in anhedonia. Of note, numerous neuroimaging and circuit physiology studies implicate the insula in the expression of anxiety and the processing of aversive states ^72, 73^. Motivated by this work, we performed an analogous analysis asking whether changes in striatal connectivity with the anterior insula area of the salience network were correlated with fluctuations in anxiety symptoms over time within each subject. In accord with our prediction, we found that striatal connectivity with anterior insula was significantly correlated with anxiety symptoms (first component score) in MDD04 and replicated this effect in MDD06 (**Fig. 4i**). Collectively, these findings show that although the salience network is stably expanded in individuals with depression, frontostriatal connectivity within this network fluctuates over time, and changes in striatal connectivity with the anterior cingulate and anterior insula track the emergence and remission of anhedonia and anxiety symptoms, respectively.

## Interpreting individual differences in salience network topology in depression

In this work, precision functional mapping in densely sampled individuals with depression revealed a marked expansion of the frontostriatal salience network that was robust and reproducible in multiple samples, with very large effect sizes relative to previously reported neuroimaging abnormalities in depression. This effect was driven primarily by network border shifts that encroached on three specific functional systems—the frontoparietal, cingulo-opercular, and default mode networks—with three distinct modes of encroachment dominating in different individuals. This effect was stable over time, insensitive to mood state, and emerged early in life in children who went on to develop depressive symptoms later in adolescence. At the same time, changes in striatal connectivity with anterior cingulate and anterior insula nodes of the salience network tracked the emergence and remission of anhedonia and anxiety, respectively, and predicted future changes in hedonic function in one individual. Of note, our analyses were enabled by large quantities of high-quality, densely-sampled multi-echo fMRI data, which may be critical for mapping individual differences in network topology precisely ^23, 74^.

Although additional work will be required to elucidate the mechanisms underlying salience network expansion in depression, key results from this report and other studies point to at least two hypotheses. First, converging evidence from multiple sources indicates that individual differences in network topology are regulated by activity-dependent mechanisms and related to the extent to which a given network is actively used. To date, most studies evaluating variability in the size of functional areas or networks across individual humans or other animals have focused primarily on the motor and visual systems. These studies have shown how different body parts have distinct representations in the primary motor cortex (M1) that differ in size, and cortical representation is closely related to the dexterity of the corresponding limb, such that the upper limbs occupy more cortical surface area than the lower limbs, as one example^75^. Motor training can increase the representation of the trained muscle or limb in M1 ^76, 77^, whereas limb amputation, casting, and congenital limb defects all decrease the representation of the disused limb and increase the representation of other body parts.^78–80^ The total surface area of primary visual cortex (V1) can vary up to three-fold in healthy young adults, and is correlated with individual differences in visual awareness^81^ and contrast sensitivity^82^. Likewise, total cortical representation of the frontoparietal network was found to be positively correlated with executive function abilities in children^26^. Together, these reports suggest that salience network expansion—accompanied by a corresponding contraction of the frontoparietal, cingulo-opercular, or default mode network—may reflect a reallocation of cortical territory and information processing priorities in individuals with depression, which could in turn contribute to alterations in salience network functions such as interoceptive awareness, reward learning, autonomic signal processing, and effort valuation.^50, 51^ This model would also predict differing deficits in functions supported by the frontoparietal, cingulo-opercular, and default mode networks in individuals expressing different modes of encroachment as in **Fig. 2g**—a hypothesis that could be tested in future studies.

Second, converging data indicate that cortical network topology is strongly influenced not only by externally modulated, activity-dependent mechanisms but also by intrinsic genetic programs^58, 83^. Numerous transcription factors regulate cell adhesion molecules, exhibit strong expression gradients across the cortical sheet during development, and covary with aspects of cortical organization, including the size or location of functional areas ^83, 84^. Deletion of these patterning factors can result in contraction or expansion of functional areas^84^. Conversely, increased expression of Emx2 increases the size of V1 and decreases the size of somatomotor areas ^85, 86^. Although our findings do not speak directly to this question, at least three observations are consistent with a role for intrinsic developmental genetic programs as opposed to exclusively activity-dependent mechanisms. First, salience network expansion was highly stable, irrespective of an individual’s current mood state, indicating that mood-related changes in network activity did not influence network size. Second, salience network expansion emerged early in life, consistent with a developmentally regulated mechanism. And third, salience network expansion was driven by spatially organized border shifts, which are known to be heritable ^5, 87^, and which tended to encroach upon neighboring networks in specific directions, expanding anteriorly and disproportionately targeting higher-order heteromodal association cortex while sparing unimodal sensorimotor areas.

## Precision mapping disentangles trait- and state-related effects in depression

Our findings may also open new avenues to addressing two fundamental challenges to using insights from clinical neuroimaging research to rethink our approach to diagnosing and treating depression. First, as noted above, MRI studies spanning two decades have identified anatomical and functional connectivity alterations that are robust and reproducible in large-scale meta-analyses but are highly variable across subjects with modest effect sizes (typically, Cohen’s *d* = 0.10 – 0.35), which complicates any efforts to glean mechanistic insights in pre-clinical models or to leverage these effects for clinical purposes. In contrast, salience network expansion was a consistent feature of a majority of individuals with depression in our sample, readily apparent on visual inspection, and associated with very large effect sizes (Cohen’s d = 1.12–1.98). Unlike other functional connectivity measures, it was also detectable without any need to correct for scanner-induced biases—which can be a significant confound for multi-site neuroimaging data ^88^. Importantly, our study was not designed to develop a neuroimaging biomarker for depression, and future work will be needed to assess the specificity of our findings with respect to other diagnoses, and evaluate its clinical utility. However, our results indicate that salience network expansion is an appealing target for biomarker development, and could have important implications for designing therapeutic neuromodulation interventions, which could have widely varying effects due to individual differences in network topology ^15, 89, 90^.

Finally, our study provides proof-of-principle data to support the use of precision functional mapping and deep, longitudinal sampling for understanding cause and effect in clinical neuroimaging studies of depression. Our analyses reveal stable, trait-like differences in salience network topology that are not only associated with depression but also emerge early in life in children with no history of depression and predict the subsequent emergence of depressive symptoms in adolescence. At the same time, they show how changes in functional connectivity strength between specific salience network nodes not only correlate with symptoms in cross-sectional samples but also track the emergence and remission of dysfunction in specific symptom domains within individuals over time, and in at least one individual, predict the future emergence of anhedonia symptoms at least one week before they occur. In this way, they show how dense sampling and longitudinal designs will open new avenues for understanding cause and effect and for designing personalized, prophylactic treatments.

---

**Extended Data Fig. 1.**
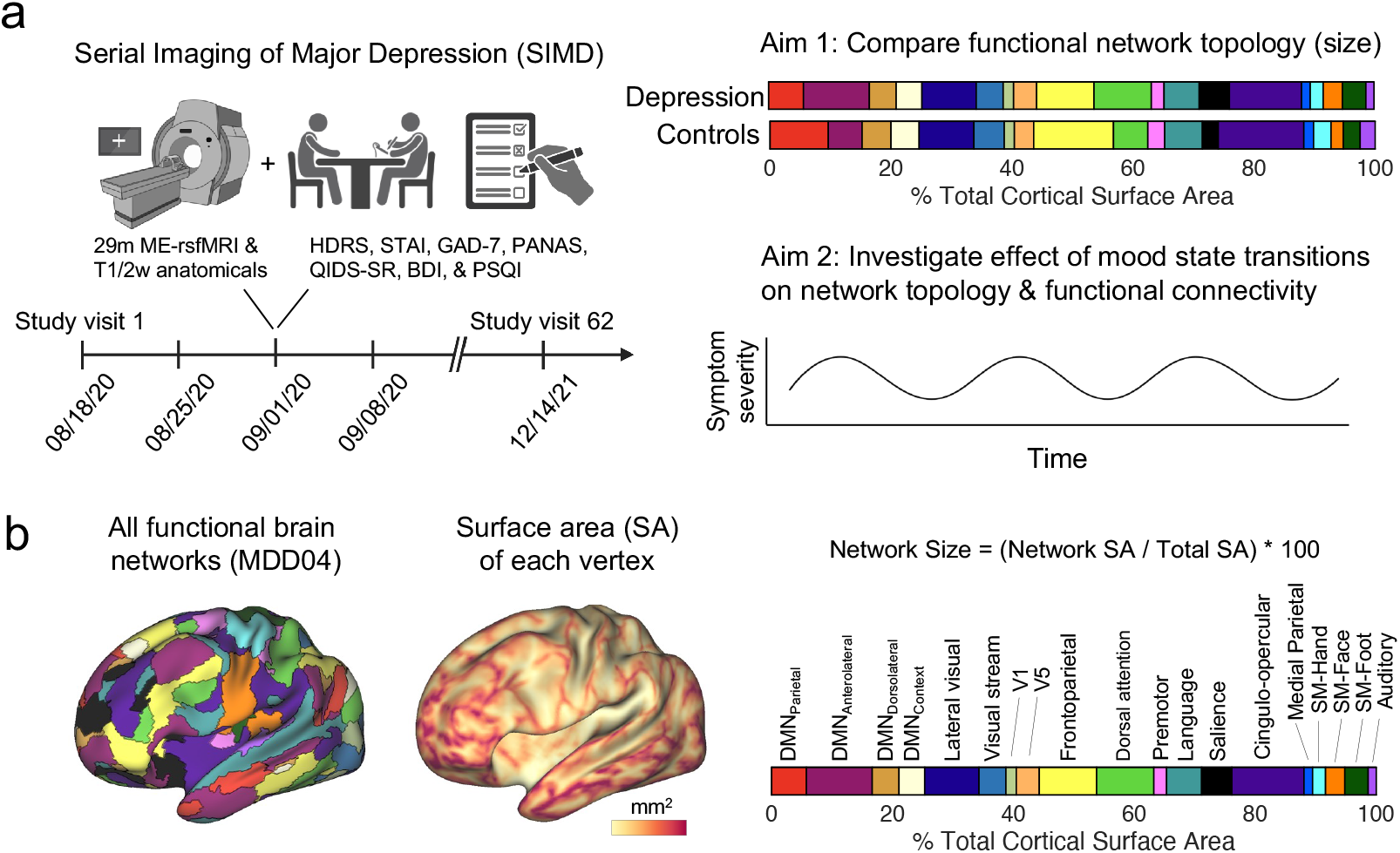
Serial Imaging of Major Depression. **a**, The SIMD project involved repeated multi-echo resting-state fMRI scans (ME-rsfMRI) and clinical assessments of six individuals with depression over long periods of time. Precision functional mapping was then used to 1) investigate differences in functional network topology, specifically size relative to healthy controls, and 2) identify which atypical aspects of network topology or connectivity are stable versus sensitive to mood state within individuals as the severity of their symptoms fluctuated, and they cycled in and out of depressive episodes. **b,** The relative contribution (size) of each functional network to the total cortical surface area was obtained by taking the total surface area of all network vertices in relation to the total cortical surface area. This approach controls for fact that each cortical vertex represents a different amount of surface area (SA). In the striatum, where each voxel represents the same amount of tissue, the relative contribution of each functional network to the total striatal volume was calculated by taking the total number of network voxels in relation to the total striatal voxels.

**Extended Data Fig. 2.**
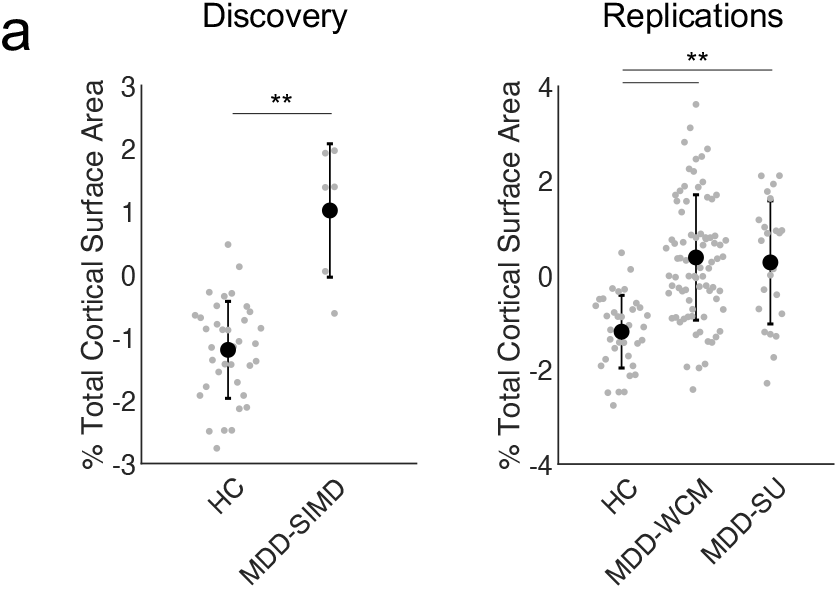
Salience network expansion remains statistically significant when controlling for sex ratio imbalance. **a**, Salience network size was regressed against sex (a variable of non-interest) and group comparisons repeated using residuals. The salience network was still significant larger in the Serial Imaging of Major Depression (MDD-SIMD) relative to healthy controls (permutation test, \*\**P* = 0.001, Bonferroni correction, *Z*-score = 6.04, Cohen’s *d* = 1.97). This effect was also replicated using data from the other imaging sites (two-tailed independent sample *t*-tests, n = 83 from Weill Cornell Medicine, MDD-WCM: *t* = 8.07, \*\**P* < 0.001, Bonferroni correction, Cohen’s *d* = 1.13; n = 23 from Stanford University, MDD-SU: *t* = 4.87, \*\**P* < 0.001, Bonferroni correction, Cohen’s *d* = 1.18).

**Extended Data Fig. 3.**
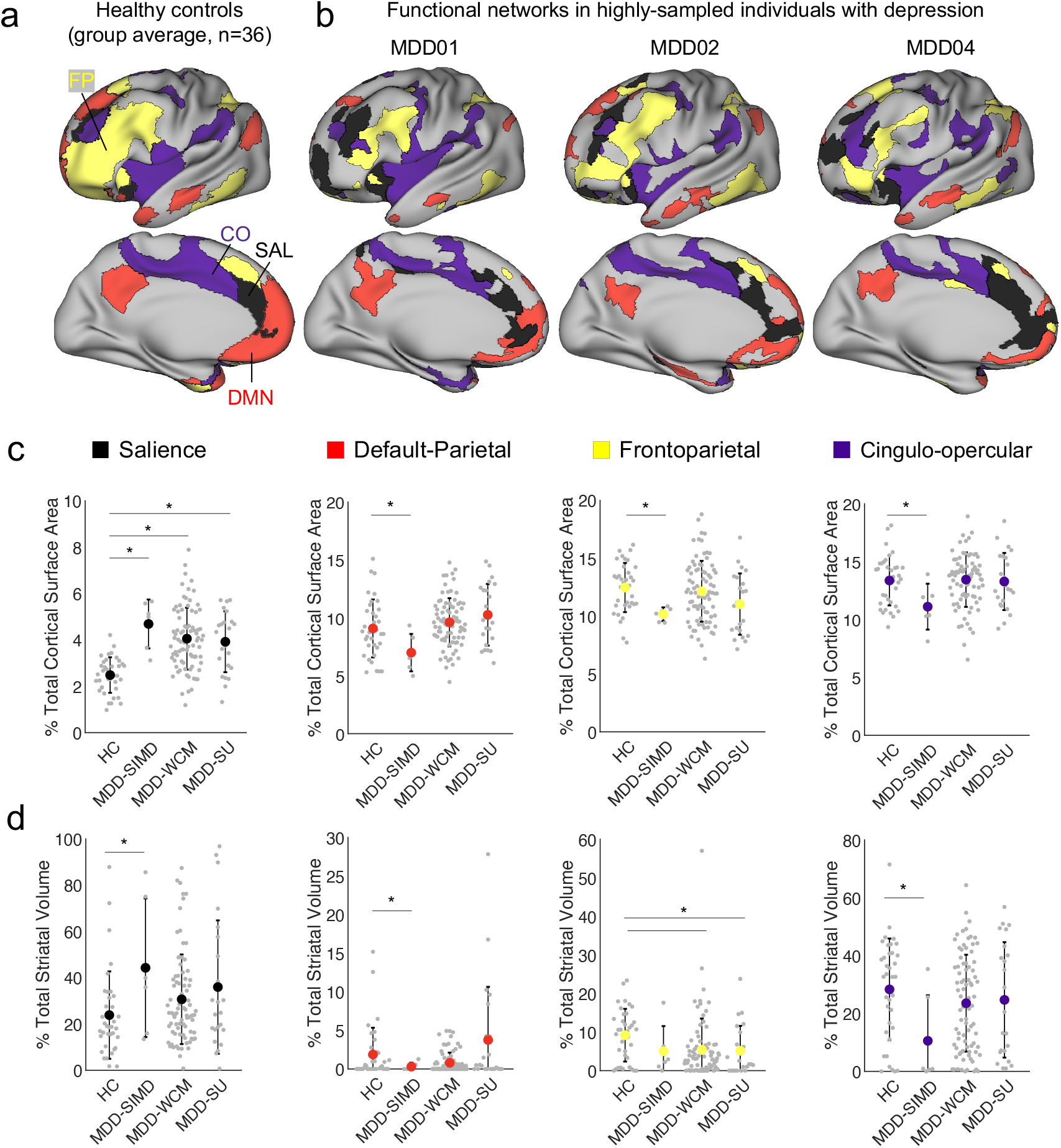
**a-b**, The salience (SAL, black), default mode (DMN, red), frontoparietal (FP, yellow), and cingulo-opercular (CO, purple) networks in a group-average map of healthy controls versus 3 representative individuals with depression. Expansion of the salience network in cortex and striatum (see **Fig. 1c-d**) was accompanied in some cases by contraction of other functional networks. **c,** In cortex, the default mode (permutation test, \**P* = 0.05, uncorrected, *Z*-score = 1.84), frontoparietal (permutation test, \**P* = 0.01, uncorrected, *Z*-score = 2.75), and cingulo-opercular (permutation test, \**P* = 0.02, uncorrected, *Z*-score = 2.45) networks were all smaller in individuals with depression in the SIMD dataset (MDD-SIMD), but this effect was not observed in either replication dataset. **d,** The striatal representation of the cingulo-opercular network was increased in the SIMD dataset relative to healthy controls (two-tailed independent sample *t*-tests, \**P* = 0.02, uncorrected, *Z*-score = 2.41), whereas the striatal representation of the frontoparietal network was decreased in the replication datasets (two-tailed paired sample *t*- tests, MDD-WCM: *t* = 2.44, \**P* = 0.02, uncorrected; MDD-SU: *t* = 2.26, \**P* = 0.03, uncorrected).

**Extended Data Fig. 4.**
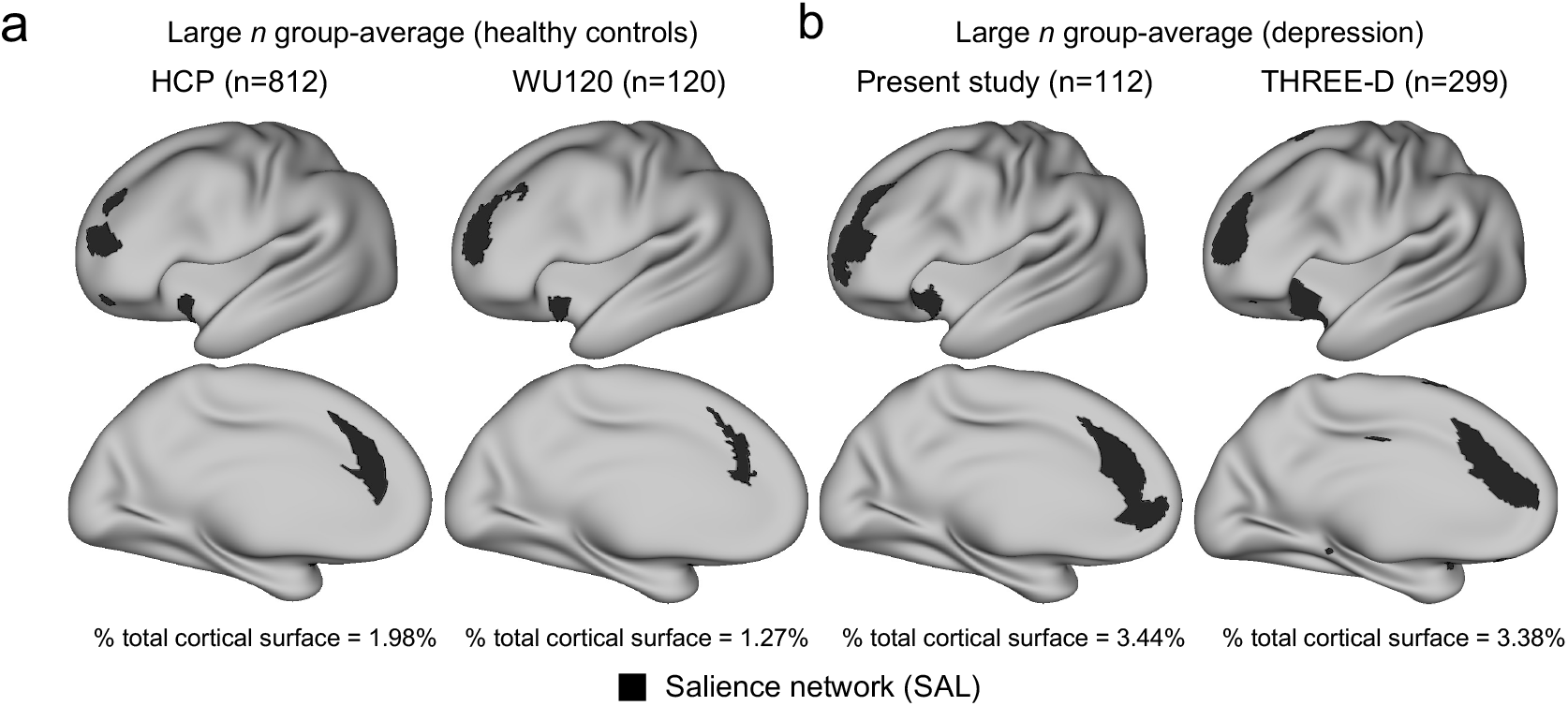
Evidence of salience network expansion in large *n* group-average datasets. **a**, Salience network mapped using two large *n* group-average data from previous studies of healthy controls occupy 1.27% and 1.98% of cortex. The group-average HCP functional connectivity matrix (which only includes subjects with resting-state fMRI data reconstructed with the r227 recon algorithm) was obtained from the S1200 release and subjected to the same precision functional mapping procedures applied to individual subjects in the main text. The WU120 salience network map was obtained directly from https://balsa.wustl.edu/jNXKl. **b**, Salience network mapped using large n group-average data and previous studies of depression occupies between 3.44% (mode assignment of all individuals with depression in current study) and 3.88% of total cortical surface area. Group-averaged functional connectivity was calculated in the THREE-D sample using group-level PCA (MELODIC Incremental Group-PCA, MIGP), and the resultant group-average FC matrix was subjected to the same precision functional mapping procedures applied to individual subjects in the main text.

**Extended Data Fig. 5.**
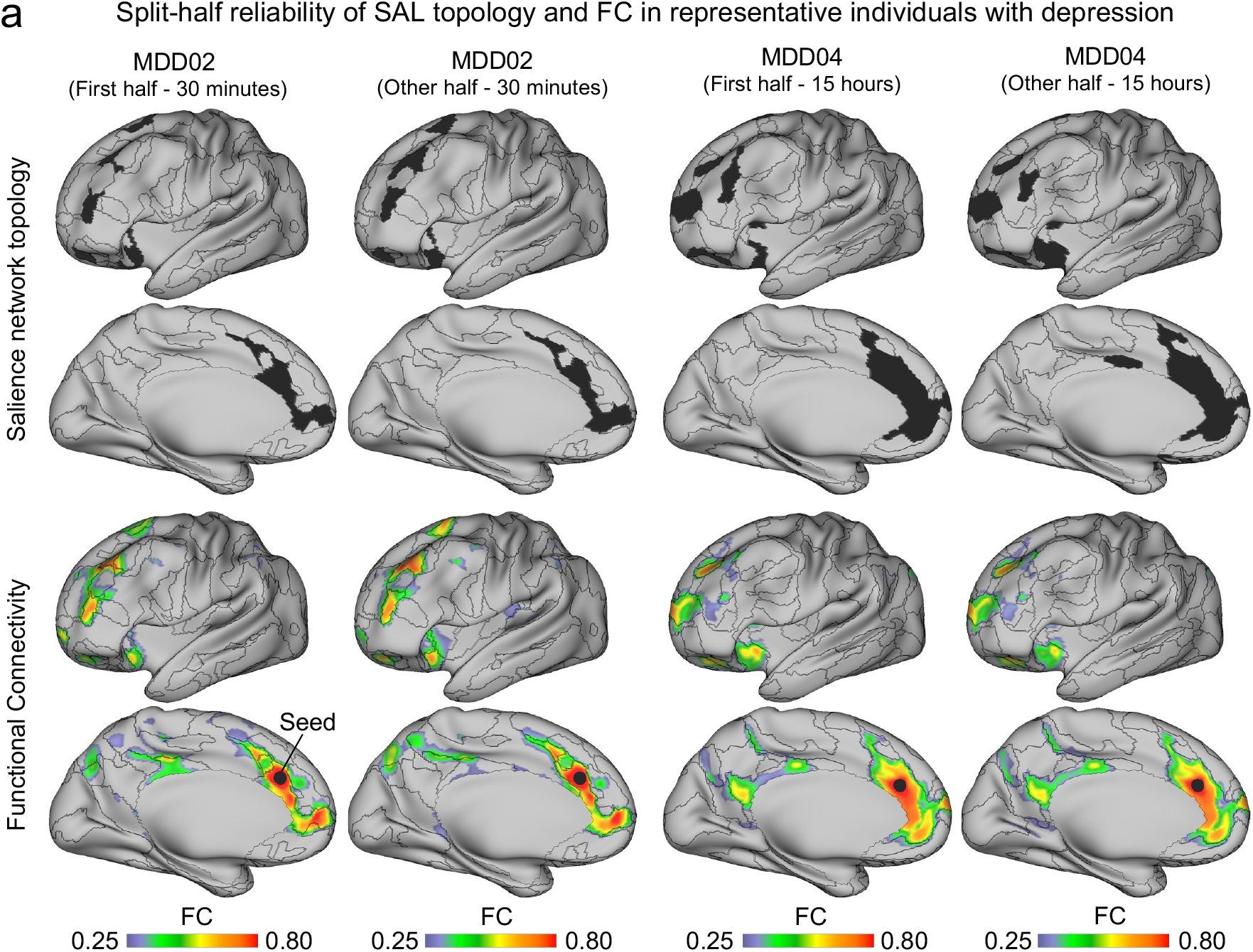
Within-person stability of salience network topology and connectivity. **a-b**, Split-half reliability testing of salience network topology and functional connectivity in the least (MDD02, 58 minutes of fMRI scanning total) and most (MDD04, 29.96 hrs. of fMRI scanning total) sampled individuals with depression from the Serial Imaging of Major Depression (SIMD) dataset.

**Extended Data Fig. 6.**
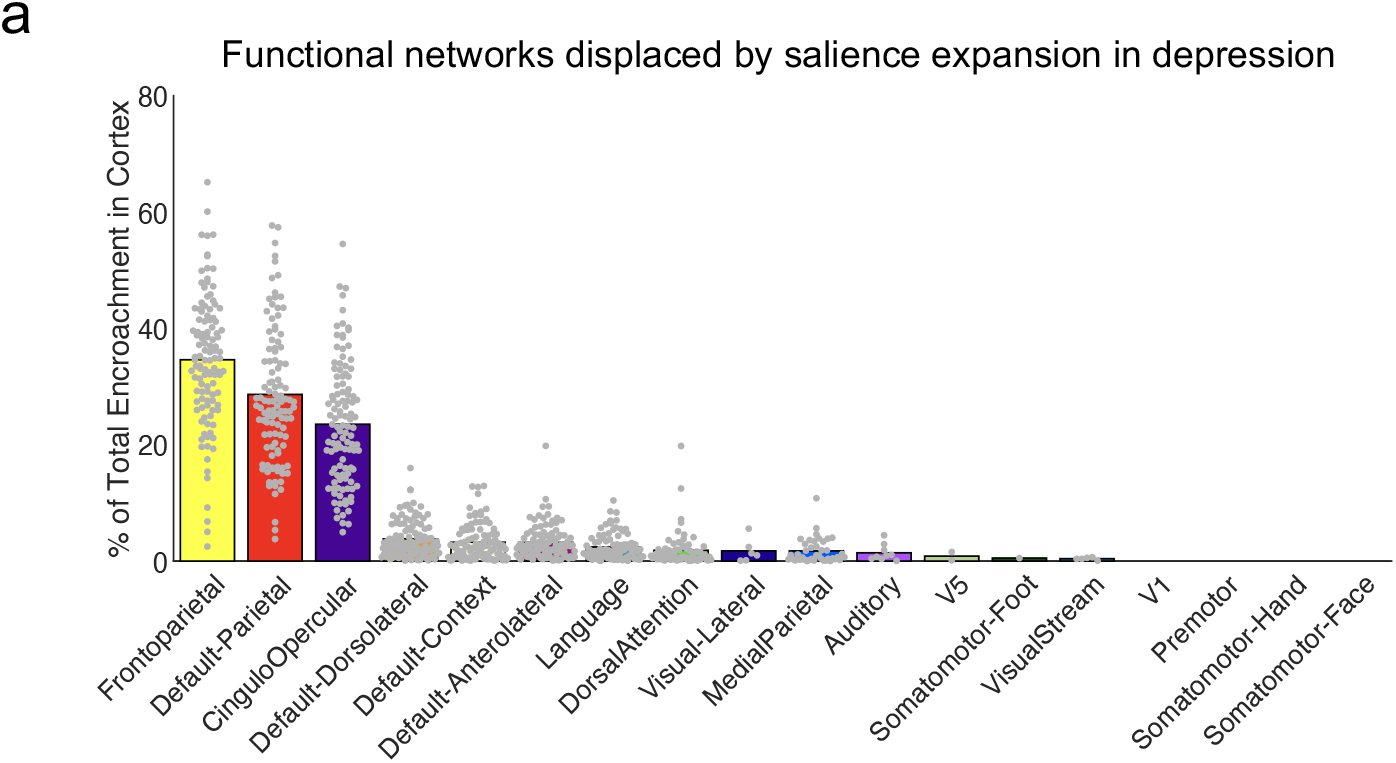
Salience network expansion in depression disproportionately affects heteromodal systems neighboring it, not unimodal sensorimotor networks. **a**, On average, nearly all (87%) of encroachment associated with salience network expansion in depression affected either the frontoparietal (34.6%), default (28.6%), or cingulo-opercular (23.5%) networks. In contrast, encroachment upon unimodal sensorimotor networks (e.g., the visual, auditory, somatomotor networks) was minimal.

**Extended Data Fig. 7.**
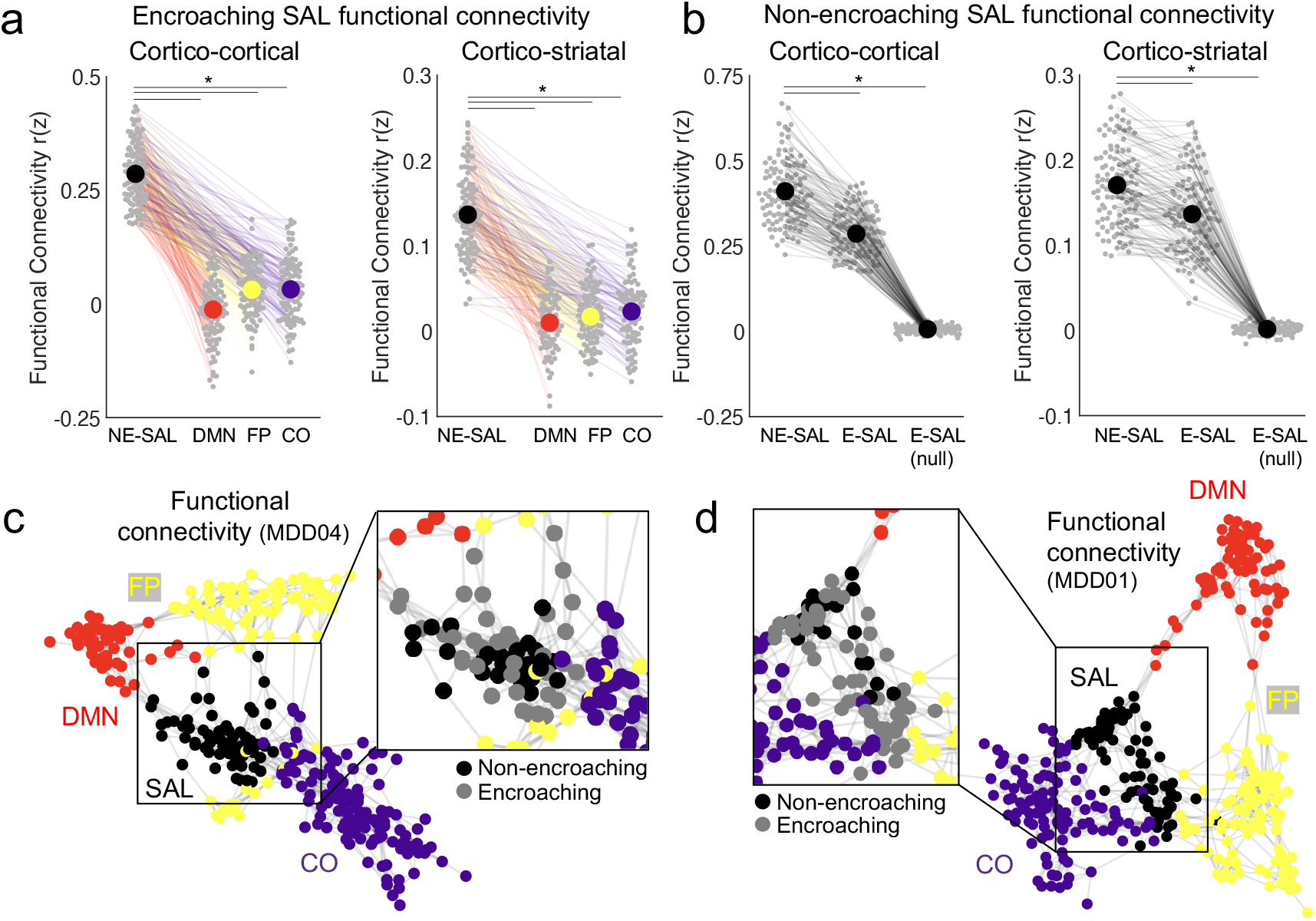
Salience network expansion in depression is associated with atypical functional connectivity. **a**, Strength of functional connectivity between encroaching nodes of the salience network, and the functional networks that typically occupy that space in healthy controls (DMN, FP, or CO). In cortex, the encroaching parts of the salience network in depressed persons were more strongly connected to the (non-encroaching, NE) parts of the salience network than with either the default mode (two-tailed t-test, *t* = 33.65, \**P* < 0.001, Bonferroni correction), frontoparietal (two-tailed *t*-test, *t* = 31.46, \**P* < 0.001, Bonferroni correction), and cingulo-opercular networks (two-tailed *t*-test, *t* = 29.18, \**P* < 0.001, Bonferroni correction). In the striatum, the encroaching parts of the salience network in depressed persons were more strongly connected to the (non-encroaching, NE) parts of the salience network than with either the default mode (two-tailed t-test, *t* = 22.37, \**P* < 0.001, Bonferroni correction), frontoparietal (two-tailed *t*-test, *t* = 22.64, \**P* < 0.001, Bonferroni correction), or cingulo-opercular networks (two-tailed *t*-test, *t* = 19.21, \**P* < 0.001, Bonferroni correction). **b,** Functional connectivity strength between non-encroaching (NE-SAL) nodes of the salience network was significantly stronger than with encroaching nodes (two-tailed *t*-tests, Cortico-cortical: *t* = 11.67, \**P* < 0.001, Bonferroni correction; Cortico-striatal: *t* = 5.54, \**P* < 0.001, Bonferroni correction) and randomly rotated versions of encroaching nodes (two-tailed *t*-test, *t* = 44.86, \**P* < 0.001, Bonferroni correction; Cortico-striatal: *t* = 38.12, \**P* < 0.001, Bonferroni correction). **c-d,** Spring graph visualizations showed how encroaching and non-encroaching nodes are more strongly connected to one another than they are with other functional networks.

**Extended Data Fig. 8.**
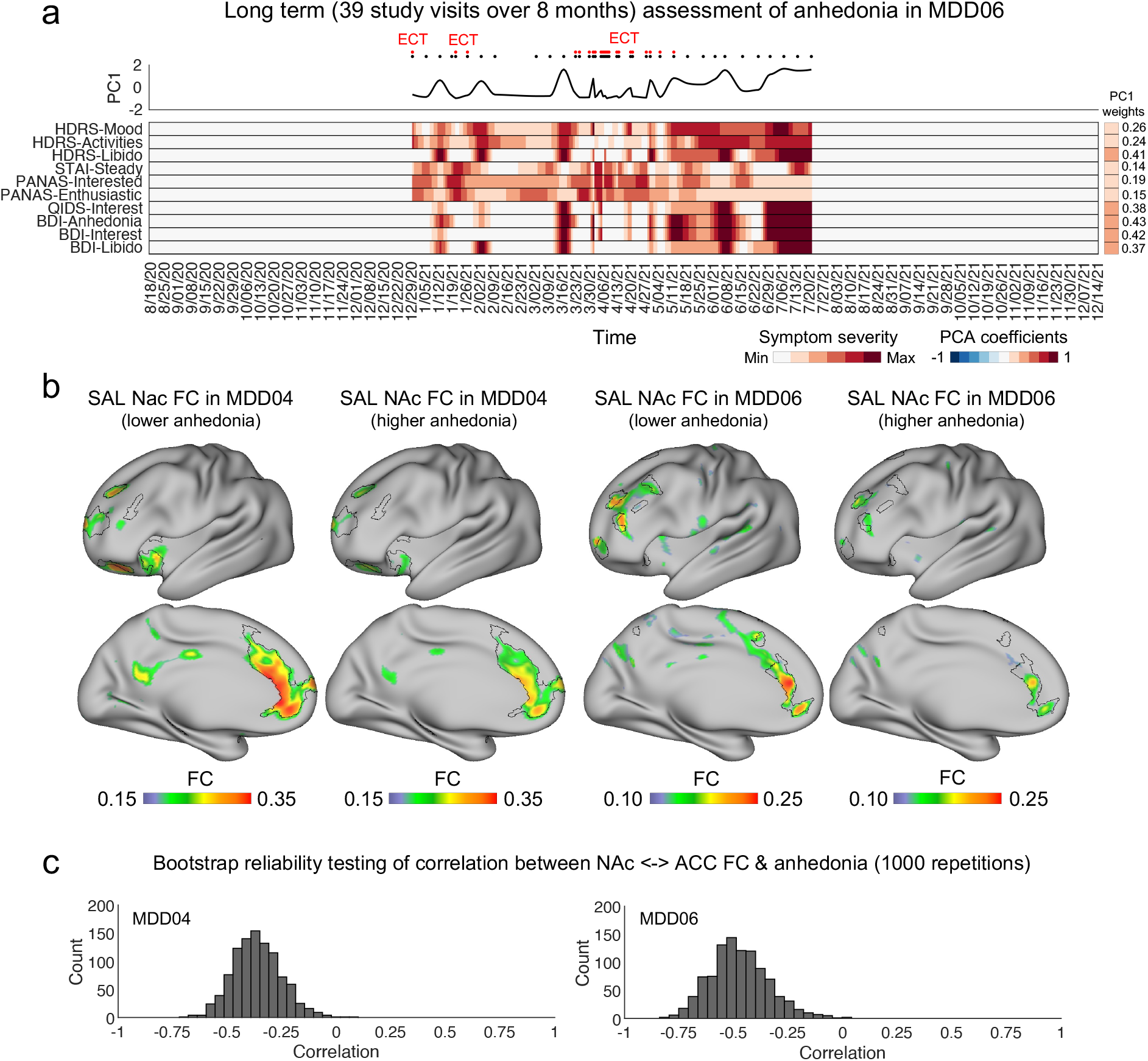
Dense-sampling of depressive symptoms and functional connectivity in a second individual with depression. **a**, A heat map summarizes fluctuations in individual items selected from a variety of clinical interviews and self-report scales related to anhedonia in an example individual (MDD06). Clinical data was resampled (using shape-preserving piecewise cubic interpolation) to days for visualization purposes (black and red dots above heat map mark the dates of study visits and ECT treatments received unrelated to the present study, respectively). **b**, Functional connectivity of salience network voxels in nucleus accumbens (NAc) when symptoms of anhedonia are low (study visits in the bottom quartile) and high (study visits in top quartile). **c**, Bootstrap resampling (iteratively selecting 50% of all time points at random, and logging correlation between nucleus accumbens ←→ anterior cingulate FC and anhedonia) indicated good stability.

## METHODS

### Datasets

The 11 datasets used in this paper are described briefly below, with additional demographic and clinical information provided in the Supplementary Information. Together, these datasets comprise more than 350 hours of fMRI data.

The depression sample collectively consisted of 112 individuals (mean age = 38.16 ± 13.81, 66F) with a diagnosis of major depression (based on DSM-IV-TR criteria and confirmed by the Mini-International Neuropsychiatric Interview administered by a trained clinician) drawn from 4 independent datasets — the Serial Imaging of Major Depression (SIMD), Weill Cornell rTMS, Stanford University rTMS, and Weill Cornell Late-onset Depression datasets.

### Serial Imaging of Major Depression (SIMD) dataset

Six individuals (mean age = 29.47 ± 8.28 years, 3F) with a primary diagnosis of major depression underwent quasi-weekly (average interval between study visits = 7.71 days ± 4.48) clinical assessments and MRI scans (2 to 124 x 14.5 minute multi-echo resting-state fMRI scans, 58 to 1,792 minutes total per-subject) longitudinally up to 1.5 years. These individuals received treatment with their psychiatrists outside of the present study (5 of the 6 individuals were stably medicated with an antidepressant medication such as an SSRI, SNRI, or Bupropion, and two were additionally taking an augmentation agent, Lithium or Lamotrigine). One individual was not taking any medications.

Each study visit began with the Hamilton Depression Rating Scale performed by a trained clinician (either study author T.N. or I.E.), followed by multiple self-report scales (State-Trait Anxiety Inventory, General Anxiety Disorder-7, Positive and Negative Affect Schedule, Quick Inventory of Depressive Symptomatology, Beck’s Depression Inventory, and The Pittsburgh Sleep Quality Index) to measure the severity of a variety of depressive symptoms.

#### Weill Cornell rTMS dataset

Eighty-three individuals (mean age = 39.10 ± 14.11 years, 47F) with a primary diagnosis of major depression that underwent clinical assessments and MRI scans (4 to 6 x 14.40 minute multi-echo resting-state fMRI scans, 57.63 to 86.45 minutes total per-subject) before and after a course of rTMS.

#### Stanford University rTMS dataset

Twenty-three individuals (mean age = 37.08± 13.41 years, 16F) with a primary diagnosis of major depression that underwent clinical assessments and MRI scans (8 x 7.28 minute multi-echo resting-state fMRI scans, 58.24 minutes total per-subject) before and after a course of rTMS.

#### Weill Cornell Late-onset Depression dataset

Five individuals (mean age = 66.60 ± 5.31 years, 5F) with a diagnosis of major depression that also met criteria for late-onset depression (age of onset at or after the age of 60) that underwent clinical assessments and MRI scans (6 x 10.64 minute multi-echo resting-state fMRI scans, 63.84 minutes total per-subject) before, during, and after a brief evidence-based psychotherapy.

The healthy control sample collectively consisted of 36 healthy adults (mean age = 31.72 ± 7.08 years, 11F) drawn from 6 independent precision functional mapping style datasets — the Weill Cornell Multi-echo ^23, 90^, MyConnectome ^12^, Midnight Scan Club (MSC) ^2^, Cast-induced Plasticity ^91^, Natural Scenes Dataset (NSD) ^92^, and Eskalibur datasets ^93^.

#### Weill Cornell Multi-Echo dataset ^23, 90^

Six individuals (mean age = 34.6 ± 9.3 years, 0F) with 0.72 to 14.69 hours of multi-echo resting-state fMRI data (2 - 61 x 14.5 minute scans).

#### MyConnectome dataset ^12^

A single 45 year-old male. This dataset was obtained from the project’s website (http://myconnectome.org/wp/data-sharing/), and includes 14 hours (84 x 10 minute scans acquired over 18 months) of preprocessed, denoised, and surface registered single-echo resting-state fMRI data. No additional preprocessing or denoising was performed.

#### Midnight Scan Club dataset ^2^

This dataset was obtained from OpenfMRI.org (https://openneuro.org/datasets/ds000224/versions/1.0.4), and includes 10 individuals (mean age = 29.1 ± 3.3 years, 5F) with 5 hours (10 x 30 minute scans acquired over two months) of preprocessed, denoised, and surface-registered single-echo resting-state fMRI data. No additional preprocessing or denoising was performed.

#### Cast-induced Plasticity dataset ^91^

This dataset was obtained from OpenfMRI.org (https://openneuro.org/datasets/ds002766/versions/3.0.0). Only one of the three individuals in this dataset was included (a single 27 year-old male, the other two subjects were already included as part of the Midnight Scan Club), and this person had 7 hours (14 x 30 minute scans acquired over consecutive days prior to the casting of their dominant upper extremity) of preprocessed, denoised, and surface-registered single-echo resting-state fMRI data. No additional preprocessing or denoising was performed.

#### Natural Scenes Dataset ^92^

This dataset was obtained in its minimally preprocessed format from Amazon Web Services (AWS), and includes 8 individuals (mean age = 26.50 ± 4.24 years, 6F) with 1.67 to 3 hours of single-echo resting-state fMRI data (22 - 40 x 6 minute scans. acquired at 7T).

#### Eskalibur dataset ^93^

This dataset was obtained in an unprocessed format from study authors S.M. and C.C, and includes 10 individuals (mean age = 31.4 ± 5.4 years, 5F) with 6.66 hours of multi-echo resting-state fMRI data (40 x 10 minute scans).

#### Adolescent Brain Cognitive Development (ABCD) dataset ^63^

The ABCD study is an ongoing longitudinal study of more than 11K 9-10-year-olds that involves annual behavioral assessments and biennial neuroimaging over ten years. We used the ABCD dataset (release 4.0) in the present study to clarify the extent to which atypical salience network topology precedes the onset of depression symptoms. Symptoms of depression in the ABCD study were operationalized using the ASEBA DSM-oriented scale for depression (“cbcl_scr_dsm5_depress_t”) from the ABCD Parent Child Behavior Checklist (CBCL). We identified n = 54 subjects (29F) meeting criteria for onset of clinical depression symptoms at the 3-year follow-up (t-score ≥ 70 at the 3-year follow-up and t-scores < 65 at previous study visits). FastTrack (unprocessed) neuroimaging data for these subjects was obtained via NDA command line utilities (https://github.com/NDAR/nda-tools). Twenty-eight subjects had sufficient data (> 40 minutes of fMRI data) for precision functional mapping.

## METHOD DETAILS

### MRI acquisition

#### Serial Imaging of Major Depression dataset

Data were acquired on a Siemens Magnetom Prisma 3T scanner at the Citigroup Biomedical Imaging Center of Weill Cornell’s medical campus using a Siemens 32-channel head coil. Two multi-echo, multi-band resting-state fMRI scans were collected using a T_2_*-weighted echo-planar sequence covering the full brain (TR: 1355 ms; TE_1_: 13.40 ms, TE_2_: 31.11 ms, TE_3_: 48.82 ms, TE_4_: 66.53 ms, and TE_5_: 84.24 ms; FOV: 216 mm; flip angle: 68° (the Ernst angle for gray matter assuming a T1 value of 1400 ms); 2.4 mm isotropic voxels; 72 slices; AP phase encoding direction; in-plane acceleration factor: 2; and multi-band acceleration factor: 6) with 640 volumes acquired per scan for a total acquisition time of 14 minutes and 27 seconds. Spin echo EPI images with opposite phase encoding directions (AP and PA) but identical geometrical parameters and echo spacing were acquired before each resting-state scan. Multi-echo T1-weighted (TR/TI: 2500/1000 ms; TE_1_: 1.7 ms, TE_2_: 3.6 ms, TE_3_: 5.5 ms, TE_4_: 7.4 ms ; FOV: 256 mm; flip angle: 8°, and 208 sagittal slices with a 0.8 mm slice thickness) and T2-weighted anatomical images (TR: 3200 ms; TE: 563 ms; FOV: 256; flip angle: 8°, and 208 sagittal slices with a 0.8 mm slice thickness) were acquired at the end of each session.

#### Weill Cornell rTMS dataset

MRI data were acquired on a Siemens Magnetom Prisma 3T machine at the Citigroup Biomedical Imaging Center of Weill Cornell’s medical campus using a Siemens 32-channel head coil. Two multi-echo, multi-band resting-state fMRI scans were collected at each study visit using a T_2_*-weighted echo-planar sequence covering the full brain (TR: 1300 ms; TE_1_: 12.60 ms, TE_2_: 29.51 ms, TE_3_: 46.42 ms, and TE_4_: 63.33 ms; FOV: 216 mm; flip angle: 67° (the Ernst angle for gray matter assuming a T1 value of 1400 ms); 2.5 mm isotropic voxels; 60 slices; AP phase encoding direction; in-plane acceleration factor: 2; and multi-band acceleration factor: 4) with 650 volumes acquired per scan for a total acquisition time of 14 minutes and 5 seconds.Spin echo EPI images with opposite phase encoding directions (AP and PA) but identical geometrical parameters and echo spacing were acquired before each resting-state scan. Multi-echo T1-weighted (TR/TI: 2500/1000 ms; TE_1_: 1.7 ms, TE_2_: 3.6 ms, TE_3_: 5.5 ms, TE_4_: 7.4 ms ; FOV: 256; flip angle: 8°, and 208 sagittal slices with a 0.8 mm slice thickness) and T2-weighted anatomical images (TR: 3200 ms; TE: 563 ms; FOV: 256; flip angle: 8°, and 208 sagittal slices with a 0.8 mm slice thickness) were acquired at the end of each session.

#### Stanford University rTMS dataset

MRI data were acquired on a GE SIGNA 3T machine at the Center for Neurobiological Imaging on Stanford University’s campus using a Nova Medical 32- channel head coil. Four multi-echo, multi-band resting-state fMRI scans were collected using a T_2_*-weighted echo-planar sequence covering the full brain (TR: 1330 ms; TE_1_: 13.7 ms, TE_2_: 31.60 ms, TE_3_: 49.50 ms, and TE_4_: 67.40 ms; flip angle: 67° (the Ernst angle for gray matter assuming a T1 value of 1400 ms); 3 mm isotropic voxels; 52 slices; AP phase encoding direction; in-plane acceleration factor: 2; and multi-band acceleration factor: 4) with 338 volumes acquired per scan for a total acquisition time of 7 minutes and 30 seconds. Spin echo EPI images with opposite phase encoding directions (AP and PA) but identical geometrical parameters and echo spacing were acquired before each resting-state scan.T1-weighted and T2-weighted anatomical images were acquired at the end of each session.

#### Weill Cornell Late-onset Depression dataset

MRI data were acquired on a Siemens Magnetom Prisma 3T machine at the Citigroup Biomedical Imaging Center of Weill Cornell’s medical campus using a Siemens 32-channel head coil. Two multi-echo, multi-band resting-state fMRI scans were collected at each study visit using a T_2_*-weighted echo-planar sequence covering the full brain (TR: 1300 ms; TE_1_: 12.60 ms, TE_2_: 29.51 ms, TE_3_: 46.42 ms, and TE_4_: 63.33 ms; FOV: 216 mm; flip angle: 67° (the Ernst angle for gray matter assuming a T1 value of 1400 ms); 2.5 mm isotropic voxels; 60 slices; AP phase encoding direction; in-plane acceleration factor: 2; and multi-band acceleration factor: 4) with 480 volumes acquired per scan for a total acquisition time of 10 minutes and 38 seconds. Spin echo EPI images with opposite phase encoding directions (AP and PA) but identical geometrical parameters and echo spacing were acquired before each resting-state scan. Multi-echo T1-weighted (TR/TI: 2500/1000 ms; TE_1_: 1.7 ms, TE_2_: 3.6 ms, TE_3_: 5.5 ms, TE_4_: 7.4 ms ; FOV: 256 mm; flip angle: 8°, and 208 sagittal slices with a 0.8 mm slice thickness) and T2-weighted anatomical images (TR: 3200 ms; TE: 563 ms; FOV: 256; flip angle: 8°, and 208 sagittal slices with a 0.8 mm slice thickness) were acquired at the end of each session.

### Anatomical preprocessing and cortical surface generation

Anatomical data were preprocessed and cortical surfaces generated using the Human Connectome Project (HCP) PreFreeSurfer, FreeSurfer, and PostFreeSurfer pipelines (version 4.3).

### Multi-echo fMRI preprocessing

Preprocessing of multi-echo data minimized spatial interpolation and volumetric smoothing while preserving the alignment of echoes. The single-band reference (SBR) images (one per echo) for each scan were averaged. The resultant average SBR images were aligned, averaged, co-registered to the ACPC aligned T1-weighted anatomical image, and simultaneously corrected for spatial distortions using FSL’s topup and epi_reg programs. Freesurfer’s bbregister algorithm ^94^ was used to refine this co-registration. For each scan, echoes were combined at each timepoint and a unique 6 DOF registration (one per volume) to the average SBR image was estimated using FSL’s MCFLIRT tool ^95^, using a 4-stage (sinc) optimization. All of these steps (co-registration to the average SBR image, ACPC alignment, and correcting for spatial distortions) were concatenated using FSL’s convertwarp tool and applied as a single spline warp to individual volumes of each echo after correcting for slice time differences using FSL’s slicetimer program. The functional images underwent a brain extraction using the co-registered brain extracted T1-weighted anatomical image as a mask and corrected for signal intensity inhomogeneities using ANT’s N4BiasFieldCorrection tool. All denoising was performed on preprocessed, ACPC-aligned images.

### Multi-echo fMRI denoising

Preprocessed multi-echo data were submitted to multi-echo ICA (ME-ICA; ^96^), which is designed to isolate spatially structured T2*- (neurobiological; “BOLD-like”) and S0-dependent (non-neurobiological; “not BOLD-like”) signals and implemented using the “tedana.py” workflow ^97^. In short, the preprocessed, ACPC-aligned echoes were first combined according to the average rate of T_2_* decay at each voxel across all time points by fitting the monoexponential decay, S(t) = S_0_e ^-t/T^^2^* . From these T_2_* values, an optimally-combined multi-echo (OC-ME) time-series was obtained by combining echoes using a weighted average (WTE = TE * e -TE/ T2*), as in ^98^. The covariance structure of all voxel time-courses was used to identify major signals in the OC-ME time-series using principal component and independent component analysis. Components were classified as either T2*-dependent (and retained) or S0-dependent (and discarded), primarily according to their decay properties across echoes. All component classifications were manually reviewed by author CJL and revised when necessary following the criteria described in ^99^. Mean gray matter time-series regression was performed to remove spatially diffuse noise. Temporal masks were generated for censoring high motion time-points using a framewise displacement (FD; ^100^) threshold of 0.3 mm and a backward difference of two TRs, for an effective sampling rate comparable to historical FD measurements (approximately 2 to 4 seconds). Prior to the FD calculation, head realignment parameters were filtered using a stopband Butterworth filter (0.2 - 0.35 Hz) to attenuate the influence of respiration ^101^ on motion parameters.

### Single-echo fMRI denoising

The following denoising procedures were applied to the NSD and ABCD datasets. Preprocessed single-echo data were submitted to ICA-AROMA ^102^. All component classifications were manually reviewed by author CJL and revised when necessary following the criteria described in ^99^. Mean gray matter time-series regression was performed to remove spatially diffuse noise. Temporal masks were generated for censoring high motion time-points, as done for the multi-echo fMRI datasets.

### Surface processing and CIFTI generation of fMRI data

The denoised fMRI time-series was mapped to the midthickness surfaces (using the “-ribbon-constrained” method), combined into the Connectivity Informatics Technology Initiative (CIFTI) format, and spatially smoothed with geodesic (for surface data) and Euclidean (for volumetric data) Gaussian kernels (σ = 2.55 mm) using Connectome Workbench command line utilities ^103^. This yielded time courses representative of the entire cortical surface, subcortex (accumbens, amygdala, caudate, hippocampus, pallidum, putamen, thalamus, brainstem), and cerebellum, but excluding non-gray matter tissue. Spurious coupling between subcortical voxels and adjacent cortical tissue was mitigated by regressing the average time-series of cortical tissue < 20 mm in Euclidean space from a subcortical voxel.

### Precision mapping of functional brain networks in individuals

A functional connectivity matrix summarizing the correlation between the time-courses of all cortical vertices and subcortical voxels across all study visits was constructed. Correlations between nodes ≤ 10 mm apart (geodesic and Euclidean space used for cortico-cortical and subcortical-cortical distance, respectively) were set to zero. Correlations between voxels belonging to subcortical structures were set to zero. Functional connectivity matrices were thresholded in such a way that they retained at least the strongest X% correlations (0.01, 0.02, 0.05, 0.1, 0.2, 0.5, 1, 2, and 5%) to each vertex and voxel and were used as inputs for the InfoMap community detection algorithm ^104^. The optimal scale for further analysis was defined as the graph threshold producing the best size-weighted average homogeneity relative to the median of the size-weighted average homogeneity calculated from randomly rotated networks, as done in ^105^. Size-weighted average homogeneity was found to be maximized relative to randomly rotated communities at the 0.1% graph density. The communities at this threshold were manually reviewed and functional network identities were assigned by study author CJL.

Functional brain networks were also mapped brain-wide using the multiplex version of the InfoMap community detection algorithm ^104, 106^. In a multiplex network, physical nodes (brain regions) can exist in multiple layers (study visits). A temporal network (node x node x study visit) summarizing the correlation between the time-courses of all cortical vertices and subcortical voxels across study visits was constructed for each patient. Correlations between nodes less than 10 mm apart (geodesic and Euclidean space used for cortico-cortical and subcortical-cortical distance, respectively) were set to zero. Correlations between voxels belonging to subcortical structures were set to zero. Links between layers were generated automatically using neighborhood flow coupling ^107^. The temporal distance between layers was constrained to 1 using the “--multilayer-relax-limit” option to encode the temporal order of study visits.

### Calculating functional network size and spatial location in individuals

We first measured the surface area (in mm^2^) that each vertex in the individual’s midthickness surface is responsible for (“wb_command --surface-vertex-areas”). Next, we calculated the relative contribution (size) of each functional network to the total cortical surface area by taking the total surface area of all network vertices in relation to the total cortical surface area. In the striatum, where each voxel represents the same amount of tissue, the relative contribution of each functional network to the total striatal volume was calculated by taking the total number of network voxels in relation to the total striatal voxels. The statistical significance of group differences in network size were evaluated using permutation tests and independent sample t-tests (the latter implemented using Matlab’s ttest2.m function). Assumptions regarding equal variance were adjusted when appropriate (based on two-sample F-tests performed using Matlab’s vartest2.m function). The relative difference between groups was calculated as the absolute difference divided by network size in healthy controls. Density maps were created by calculating the percentage of individuals with salience network representation at each cortical vertex or striatum voxel.

### Classification analysis

A support vector machine classifier, where class labels were diagnosis status (healthy control or depression) and features were functional brain network size in each individual, was trained using leave-one-out cross-validation. Specifically, a classifier was fit using all of the data except for one subject, and a prediction regarding that held-out individual’s diagnosis was made using Matlab’s fitcsvm.m and predict.m functions, respectively. Classification accuracy was calculated as the percentage of correct predictions, and statistical significance assessed using permutation tests. Confusion matrix was created using Matlab’s confmat.m function. Feature importance was evaluated by iteratively omitting each functional network and calculating the resulting loss in accuracy. The Synthetic Minority Oversampling Technique (SMOTE ^57^) was used to prevent classification bias in favor of the majority class, and was performed on training data only to prevent data leakage.

### Evaluating how salience network expansion displaces other functional systems

The parts of each depressed individual’s salience network map that did and did not overlap with the salience network in the group average healthy control map were operationalized as “non-encroaching” and “encroaching”, respectively. Encroaching clusters were identified (“wb_command -cifti-find-clusters”) and were classified as border shifts if any part of the cluster was within 3.5 mm (in geodesic space) of a salience network vertex in the group average healthy control map, and as ectopic intrusions if they did not, as done in ^5^.

### Longitudinal analyses relating changes in connectivity with symptom severity

Three separate clinicians (study authors I.E, J.D.P, N.S.) independently selected items they thought captured anhedonia-like symptoms from the battery of clinical scales administered to the SIMD subjects at the start of each study visit. The consensus items were min-max normalized, adjusted for valence (so that higher scores index worse symptoms across all items), and subjected to a principal component analysis to extract an overall score (PC1) of anhedonia at each study visit. Functional connectivity strength between all pairs of cortical and striatal nodes of the salience network was calculated for each study visit, separately, and correlated with anhedonia PC1. Correlations not exceeding chance (based on null distribution of correlation coefficients obtained using shuffled clinical data) were set to zero. Circular permutation tests (using Matlab’s circshift.m function) were used to preserve temporal autocorrelation. Cross-correlation analyses were performed using Matlab’s crosscorr.m function (with “NumLags” set to 2). For the cross-sectional analysis, the overall anhedonia score was calculated using baseline clinical data.

### Data availability

Data from the Weill Cornell Multi-echo and Eskalibur datasets are available on reasonable request from C.J.L, I.E., J.D.P, C.L., and S.M., C.C. Data from the MyConnectome dataset is available from the project’s website (http://myconnectome.org/wp/data-sharing/) and in the openneuro repository at https://openneuro.org/datasets/ds000031/versions/2.0.2. Data from the Midnight Scan Club dataset is available in the openneuro repository at https://openneuro.org/datasets/ds000224/versions/1.0.4. Data from the Cast-induced plasticity dataset is available in the OpenNeuro repository at https://openneuro.org/datasets/ds002766/versions/3.0.0. Data from the Natural Scenes Dataset is available from Amazon Web Services (AWS) at https://registry.opendata.aws/nsd/. The ABCD data used in this report are from Annual Release 4.0. Data from individual subjects with depression are a part of ongoing clinical trials and not publicly available at this time.

### Code availability

Code for preprocessing multi-echo fMRI data is maintained in an online repository (https://github.com/cjl2007/Liston-Laboratory-MultiEchofMRI-Pipeline). Code for performing precision functional mapping and code specific to the analyses performed in this manuscript are maintained in an online repository (https://github.com/cjl2007/PFM-Depression). Software packages incorporated into the above pipelines for data analysis included: Matlab R2019a, https://www.mathworks.com/; Connectome Workbench 1.4.2, http://www.humanconnectome.org/software/connectome-workbench.html; Freesurfer v6, https://surfer.nmr.mgh.harvard.edu/; FSL 6.0, https://fsl.fmrib.ox.ac.uk/fsl/fslwiki; and Infomap, https://www.mapequation.org.

## Acknowledgements

We thank the staff at the Citigroup Biomedical Imaging Center for assistance with data collection. This work was supported by grants to C.L. from the National Institute of Mental Health, the National Institute on Drug Addiction, the Hope for Depression Research Foundation, and the Foundation for OCD Research. C.J.L. was supported by an NIMH F32 National Research Service Award (F32MH120989). N.S. was supported by K23 MH123864. Work on "Personalized Therapeutic Neuromodulation for Anhedonic Depression" is supported by Wellcome Leap as part of the Multi-Channel Psych Program.

## Author contributions

Conceptualization, C.J.L, I.E., T.N., J.D.P., C.L.; Methodology, C.J.L, I.E., E.M.G., J.D.P., L.G., Investigation, C.J.L, I.E.,T.N., S.Z., A.Z., N.M., M.J., A.A., E.G., D.W.; Funding Acquisition, N.W., K.K., C.C.G., F.M.G., C.L.; Resources, J.D.P, B.Z., F.V., Z.G.D, D.M.B, C.L.; Writing – Original Draft, C.J.L., I.E., J.D.P., C.L.; Writing – Review & Editing, C.J.L., I.E., J.D.P., E.M.G., K.K., A.A., S.M., N.S., E.G., L.G., F.V., Z.G.D, D.M.B, C.L.; Supervision, F.M.G., L.V., N.S., C.L.

## Ethics declarations

C.L. and C.J.L are listed as inventors for Cornell University patent applications on neuroimaging biomarkers for depression that are pending or in preparation. C.L. has served as a scientific advisor or consultant to Compass Pathways PLC, Delix Therapeutics, and Brainify.AI. The other authors declare no competing interests.

## Clinical trial information

A portion of the data used in this report was obtained from the Biomarker-guided rTMS for Treatment Resistant Depression study (NCT04041479), Randomized Controlled Trial of Conventional vs Theta Burst rTMS (HFL vs TBS) (NCT01887782), and Accelerated TMS for Depression and OCD studies (NCT04982757).

